# Mechano-induced homotypic patterned domain formation by monocytes

**DOI:** 10.1101/2023.07.27.550819

**Authors:** Wenxuan Du, Jingyi Zhu, Yufei Wu, Ashley L. Kiemen, Sean X. Sun, Denis Wirtz

## Abstract

Matrix stiffness and corresponding mechano-signaling play indispensable roles in cellular phenotypes and functions. How tissue stiffness influences the behavior of monocytes, a major circulating leukocyte of the innate system, and how it may promote the emergence of collective cell behavior is less understood. Here, using tunable collagen-coated hydrogels of physiological stiffness, we show that human primary monocytes undergo a dynamic local phase separation to form highly patterned multicellular multi-layered domains on soft matrix. Local activation of the β2 integrin initiates inter-cellular adhesion, while global soluble inhibitory factors maintain the steady-state domain pattern over days. Patterned domain formation generated by monocytes is unique among other key immune cells, including macrophages, B cells, T cells, and NK cells. While inhibiting their phagocytic capability, domain formation promotes monocytes’ survival. We develop a computational model based on the Cahn-Hilliard equation, which includes combined local activation and global inhibition mechanisms of intercellular adhesion suggested by our experiments, and provides experimentally validated predictions of the role of seeding density and both chemotactic and random cell migration on pattern formation.

## Introduction

The human body is composed of tissues that span a wide range of stiffnesses^1^. Human tissues can be as soft as 11 Pa (intestinal mucus) and as stiff as 20 GPa (cortical bone)^2^. Tissue stiffness is also altered during aging^3–5^ and under various disease conditions such as cancer^6–9^ and inflammation^10^. Mesenchymal and epithelial cells have evolved complex molecular mechanisms to sense and respond to these different environmental mechanical cues to differentiate, signal, and migrate. How immune cells respond to mechanical cues has received significantly less attention. In particular whether and how monocytes, which are pivotal components in the innate immune response, respond to microenvironments of different stiffness is not well understood^11^. Microenvironmental molecular signals can differentiate monocytes into monocyte-derived macrophages and dendritic cells, which enables them to orchestrate innate and adaptative immune responses^12, 13^. Upon infection, inflammation or tumorigenesis, classic monocytes (CD14^+^CD16^-^) that mature in the bone marrow and emigrate to the peripheral blood constantly traffic between blood vessels and soft tissues^14^, making them more likely to encounter complex micro-environments of different stiffness.

Imaging of pre-cancer and tumor tissues has revealed spatially heterogeneous immune cell hot spots – dense aggregates of immune cells – in the stomal region of precursor lesions, such as PanIN in the pancreas, as a reservoir for future infiltration^15, 16^. Accumulating studies also suggest that homotypic immune cell aggregation is a physiologically relevant process that plays a vital role in pathogen clearance and cancer metastasis for B cells^17, 18^, neutrophils^19, 20^, dendritic cells^21^ and monocytes/macrophages^22^. Cells connect with neighboring cells and their milieu by forming molecular links driven by adhesion molecules, including cadherins, selectins, integrins, Ig-like adhesion molecules, and mucins^23^. Integrins are highly expressed on the surface of monocytes, which have been found to be essential for the tethering and rolling process of monocyte extravasation and as one of the major mediators for cell-matrix adhesion via focal adhesion complex^24–26^. Whether and how matrix mechanical properties promote or modulate cell aggregation is unclear.

The observed homotypic immune cell aggregates are reminiscent of dynamic phases observed in active matter systems. In particular, flocking transitions seen during collective movement of fish and birds also produce high density aggregates^27, 28^. More recent theoretical work has revealed rich phase behavior of self-propelled particles in terms of the underlying microscopic interactions^29, 30^. Different from flocking, which are governed by local short-range interactions between self-driven particles, cells possess both short range interaction and long-range signals, and therefore potentially can generate complex behavior. Cells can respond to differences in matrix stiffness (mechanosensing), secrete factors, thrive to survive and proliferate, and move along local chemotactic gradients. Moreover, cell movement is also distinct from propelled particles, and are instead described by persistent random walks (PRW)^31, 32^. These aspects lead us to hypothesize that immune cells interacting in a population would reveal new dynamic phases and behaviors.

In this study, we find that physiological and pathological matrix stiffness can spontaneously trigger monocyte homotypic domain formation via the over-expression of β2 integrin to promote long term cell viability. We propose a phenomenological model based on Cahn-Hilliard equation^33, 34^ which incorporates activation of β2 integrins as a local activator and self-inhibitory soluble factors secreted by aggregated monocytes as global inhibitors, which simulate the patterned domain formation observed on collagen-coated hydrogels. Effects of cell seeding density and cell chemotaxis *vs*. random migration on monocyte homotypic domain patterns are predicted by computational simulations and validated in corresponding experiments.

## Results

### Soft matrix induces multicellular domain formation of monocytes

Figure 1A shows a localized high density of leukocytes in human pancreas tissue. Two consecutive 5-μm thick tissue sections were respectively stained with hematoxylin and eosin to enable visualization of the pancreatic microanatomy (top) and immuno-cytochemically stained for the cell-surface antigen CD45 to label all leukocytes (bottom). The function and mechanism of such leukocyte-rich domains remain unclear. To investigate the minimum necessary conditions for leukocytes to form such aggregates *in vitro* and assess the potential role of matrix stiffness, we placed freshly isolated human classical (CD14^+^CD16^-^) monocytes on collagen-I coated polyacrylamide gel substrates of stiffness 0.5 and 100 kPa, as well as standard cell culture plastic substrates (∼5 GPa). These substrates were chosen to mimic the range of tissue stiffness encountered by monocytes during disease progression of soft tissue such as the transition from the relatively soft microenvironment of the normal human breast to the comparatively stiff microenvironment of breast cancer^35^.

**Fig. 1.**
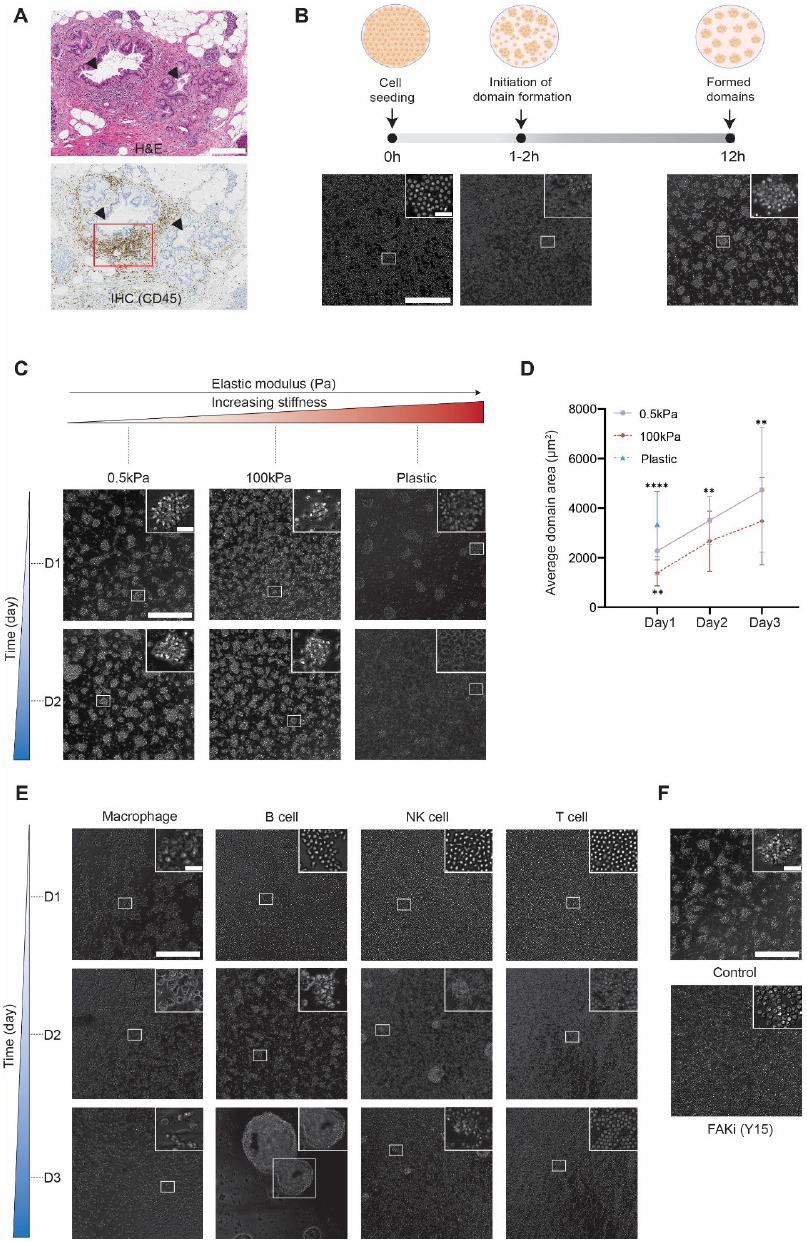
Mechano-induced homotypic domain formation of human primary monocytes on substrates of different stiffness. (**A**) Representative sections stained with hematoxylin and eosin (top, H&E) and CD45 (bottom, leukocyte common antigen) showing an immune hot spot in a human pancreatic tissue containing a precursor lesion (PanIN). The large gland on the left is ‘hot’ with aggressive accumulation of leukocytes (highlighted in red box) while the small gland on the right is relatively ‘cold’ with minimal immune cell infiltration. Scale bar, 300 μm. (**B**) Freshly isolated human primary classical (CD14^+^CD16^-^) monocytes formed compact multicellular domains when placed on a 0.5 kPa collagen I-coated substrate. Representative phase-contrast images were obtained with a 20X objective at 0h, 2h, and 12h. Monocytes spontaneously form pre-domain spots around 1-2h after initial seeding. After 12h incubation, these spots progress to highly patterned domains (see Supplementary Video 1 for domain formation process). (**C**) Representative phase-contrast images of monocytic domain formation on (soft) 0.5 kPa, (stiff) 100 kPa and (hard) plastic collagen I-coated substrates obtained via a 20X objective on day 1 and day 2. (**D**) Average areas of domains formed by monocytes on substrates of different stiffness. Monocytic domains did not form on collagen-coated plastic substrates and were therefore not characterized on day 2 and day 3. (**E**) Representative phase-contrast images of macrophages / B cell / NK cell / T cell domain formations on 0.5 kPa soft substrate obtained on day 1, day 2 and day 3. None of the lymphocytes initiated homotypic aggregation for the first 24h like monocytes’ quick response to mechanical cues of the substrate. Macrophages, however, displayed homotypic aggregation on day 1 but failed to maintain. Representative day 1 images were taken with 10x images while day 2/3 images were taken with 20x objective. (**F**) Inhibition of focal adhesion kinase using 20 μM Y15 abrogated domain formation. Representative 20x images were taken on day 1. Scale bars, 300 μm (insets, scale bar, 30 μm). Unpaired student’s t-test was used for statistical analysis. Results are presented as mean ± standard deviation.

Following seeding as a monolayer on 0.5 kPa collagen-I coated substrates, we observed that monocytes remained mostly featureless for an “incubation” time of 1-2 h, before a rapid phase transition occurred and multicellular aggregates formed (Supplementary Video 1). This incubation time for domain formation, during which we could observe no hint of phase separation, was remarkably consistent across donors and across different wells for a given donor (Supplementary Fig. 1B). In the first few hours post-transition, the borders of these aggregates were relatively diffuse. After ∼12h, these aggregates annealed and formed well-defined spatially patterned muti-layered aggregates, referred below as “domains” (Fig. 1B). In contrast, domain formation of monocytes placed on 100 kPa (stiff) substrates experienced a significant delay, resulting in the development of irregular domains on day 1 (Fig. 1C). Monocytes did not show long-lasting domains on collagen-I coated plastic culture dishes (∼ 5GPa). Monocytes placed on this hard substrate formed highly transient domains, whose average size was significantly larger compared to those on the 0.5 kPa and 100 kPa substrate on day 1 (Fig. 1D). But these domains collapsed after day 2 (Fig. 1C). Additionally, enlarged domain areas were observed on 0.5 kPa and 100 kPa substrates from day 1 to day3, which were potentially caused by joining of single monocytes into established domains or “flatting” of the multi-layered domain structure.

We asked if the above phenomenon was universal among immune cells or specific to monocytes. We placed either primary human monocyte-derived macrophages, B cells, NK cells or T cells on 0.5 kPa substrates and assessed potential phase separation. Macrophages initiated instant homotypic aggregation after seeding (Supplementary Video 2) but formed continuous cell clusters instead of spatially patterned domains, which is potentially driven by the over-expression of adhesion molecules during differentiation. Moreover, the majority of macrophage aggregates underwent disintegration on day 2 and single cells with clear mesenchymal morphology were observed strongly attached to the substrate on day 3 (Fig. 1E). Surprisingly, none of these common lymphocytes were capable of initiating domain formation when exposed to mechanical cues of the soft substrate, suggesting the unique ability of monocytes to form patterned aggregates. T cells remained a perfect monolayer for three consecutive days, while NK cells only formed small, sparsely distributed, diffuse clusters. For B cells which are known for aggregation after stimulation^17^ and chemotactic collective cell motility^18^, we observed patterned domain formation after 48h, but these domains subsequently merged, suggesting a macroscopic phase separation, and even partially detached from their collagen-I coated substrate (Fig. 1E).

To further support the hypothesis that monocyte domain formation was induced via mechano-sensing, we treated the monocytes with an inhibitor of the major mechano-sensing protein focal adhesion kinase (FAK)^36–38^. Monocytes did not exhibit any evidence of domain formation (Fig. 1F, Supplementary Video 3). This validates that mechano-sensing plays an indispensable role in the process of domain formation by monocytes. Together these results suggest that monocytes can spontaneously form regular multicellular domains, a process greatly influenced by the stiffness of the underlying matrix.

### Phase separation promotes cell survival and inhibits phagocytosis

Next, we investigated possible functional outcomes of monocyte domain formation. Monocytes seeded on 0.5 kPa collagen-coated substrate showed significantly higher viability than monocytes seeded on collagen-coated plastic and glass substrates within a time frame of 72h, for which collapsed domains and cellular apoptosis were clearly observed at days 2 and 3 (Fig. 2A). Quantitative live cell viability assay via PrestoBlue showed a dramatic decrease in signal from monocytes on plastic and glass substrates in contrast to cells on 0.5 kPa substrate. This result indicates an underlying functional connection between the ability of monocytes to form and maintain domains and their viability (Fig. 2B). Monocytes were also harvested after 72h from different substrates and analyzed using flow cytometry. While debris remained at a relatively low value of 10.7% for monocytes harvested from 0.5 kPa substrate, elevated proportions of debris were observed on harder substrates, 25.1% for plastic and 77.1% for glass (Fig. 2C). Finally, more propidium iodide positive monocytes were observed through immunofluorescence imaging and flow cytometry, which worked as extra validations of reduced cell viability on non-physiological substrates (Fig. 2C).

**Fig. 2.**
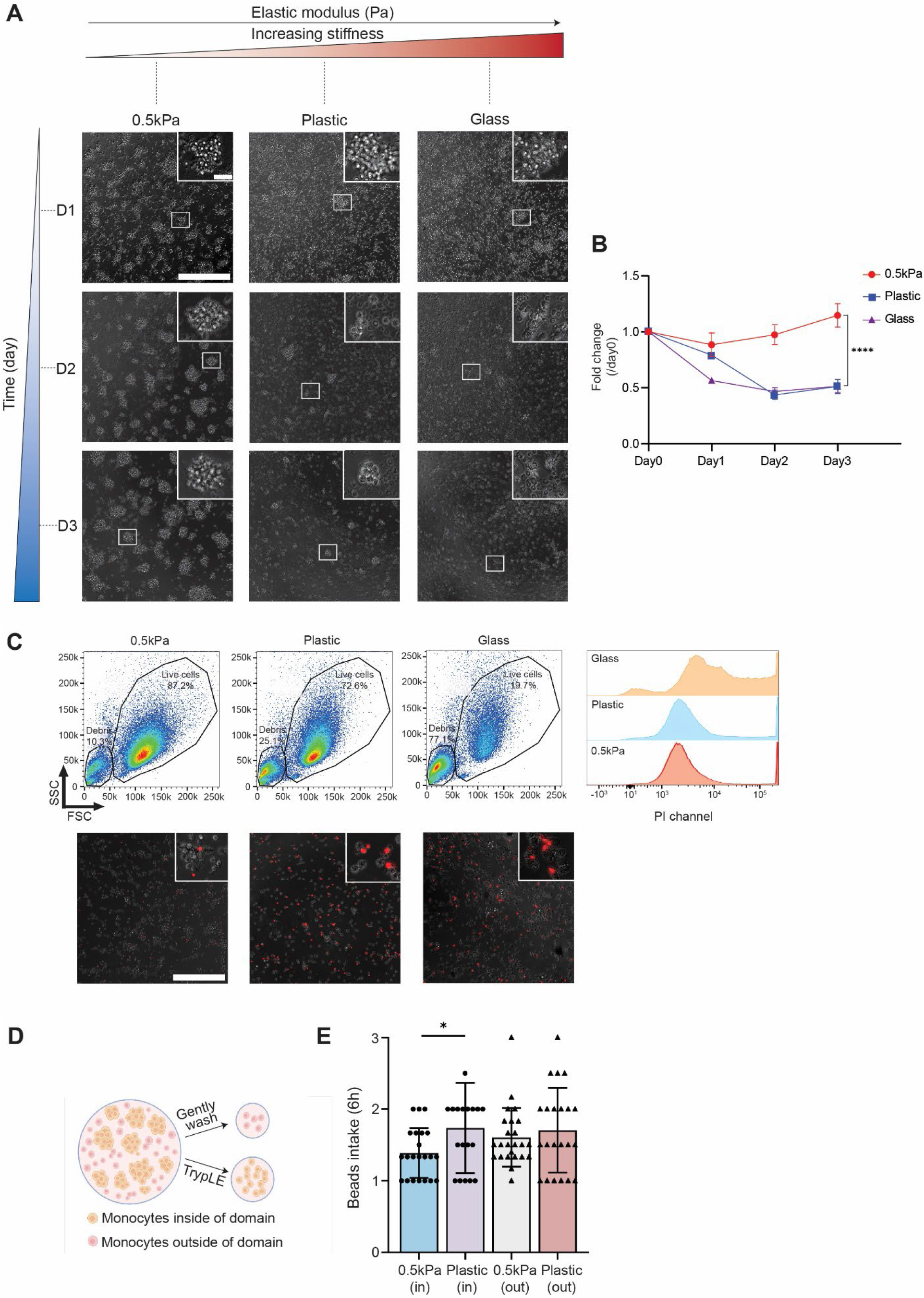
Effects of homotypic domain formation on monocyte viability and phagocytosis. (**A**) Domain formation and long-term maintenance on collagen I-coated 0.5 kPa, plastic and glass substrates. Representative phase-contrast images were taken with a 20X objective on days 1, 2 and 3. (**B**) Fold change in PrestoBlue signal normalized to day 0 when seeded. Significant cell death was observed for monocytes seeded on plastic and glass substrate with dramatically decrease of PretoBlue signals compared to day 0. In the meantime, monocytes seeded on 0.5 kPa substrate maintained a similar level of metabolic activity during the 3-day monitoring window. Flow cytometry results of monocytes harvested from 0.5kPa, plastic and glass substrates on day 3. respectively. More cell debris and propidium iodide positive cells (died cells) were observed when monocytes were seeded on plastic and glass substrate. Representative Immunofluorescence images taken with 20x objective on different substrates were shown via merging phase contrast channel and PI channel. (**D**) Illustration cartoon that distinguishes monocytes inside and outside domains. (**E**) Characterization of phagocytosis capabilities of monocytes inside and outside of domains, measured via quantification of GFP-positive Carboxylate-modified polystyrene beads (1 μm) intake after 6 hours. Scale bars = 300 μm (insets, scale bar = 30 μm). Unpaired student’s t-test was used for statistical analysis. Results are presented as mean ± standard deviation.

To study whether substrate stiffness and domain formation had any impact on the phagocytic ability of monocytes, we evaluated the number of internalized latex beads. We distinguished monocytes located within the domains as “insiders” and those outside the domains as “outsiders” (as illustrated in Fig. 2D). On the 0.5 kPa substrate, insiders and outsiders exhibited similar levels of bead phagocytosis, with insiders averaging 1.6±0.4 particles per cell and outsiders averaging 1.4±0.3 particles per cell. Likewise, on the plastic substrate, there was no significant difference in phagocytosis ability between insiders and outsiders (Fig. 2D). However, “insiders” on 0.5 kPa substrate displayed a small but statistically significant lower phagocytic activity compared to those on plastic substrate, which suggests a negative correlation between monocyte viability and phagocytosis.

### Matrix stiffness modulates β2 integrin expression, which mediates homotypic aggregation

Because focal adhesions are integrin-containing structures involved in the crosstalk between matrix and intracellular actin networks, we hypothesized that a specific subtype of integrins mediated domain formation. We compared the expressions of cell surface integrins for monocytes collected from 0.5 kPa substrate at different time points. Among tested integrins, we found that integrin α4, αM, and β2 exhibited increased expression by day 3 (for which we observed well-defined stabilized domain patterns) compared to day 1 (onset of domain formation), suggesting their potential roles in initiating and maintaining the formation of these domains (Fig. 3A). To validate these findings, we treated monocytes seeded in monolayer form (at the initialization stage of domain formation) with inhibitory antibodies targeting these three upregulated integrins. Only the anti-β2 integrin antibody completely abrogated domain formation (Fig. 3B). Moreover, the inhibitory anti-β2 integrin antibody successfully dispersed pre-formed domains (Fig. 3C, Supplementary Video 4).

**Fig. 3.**
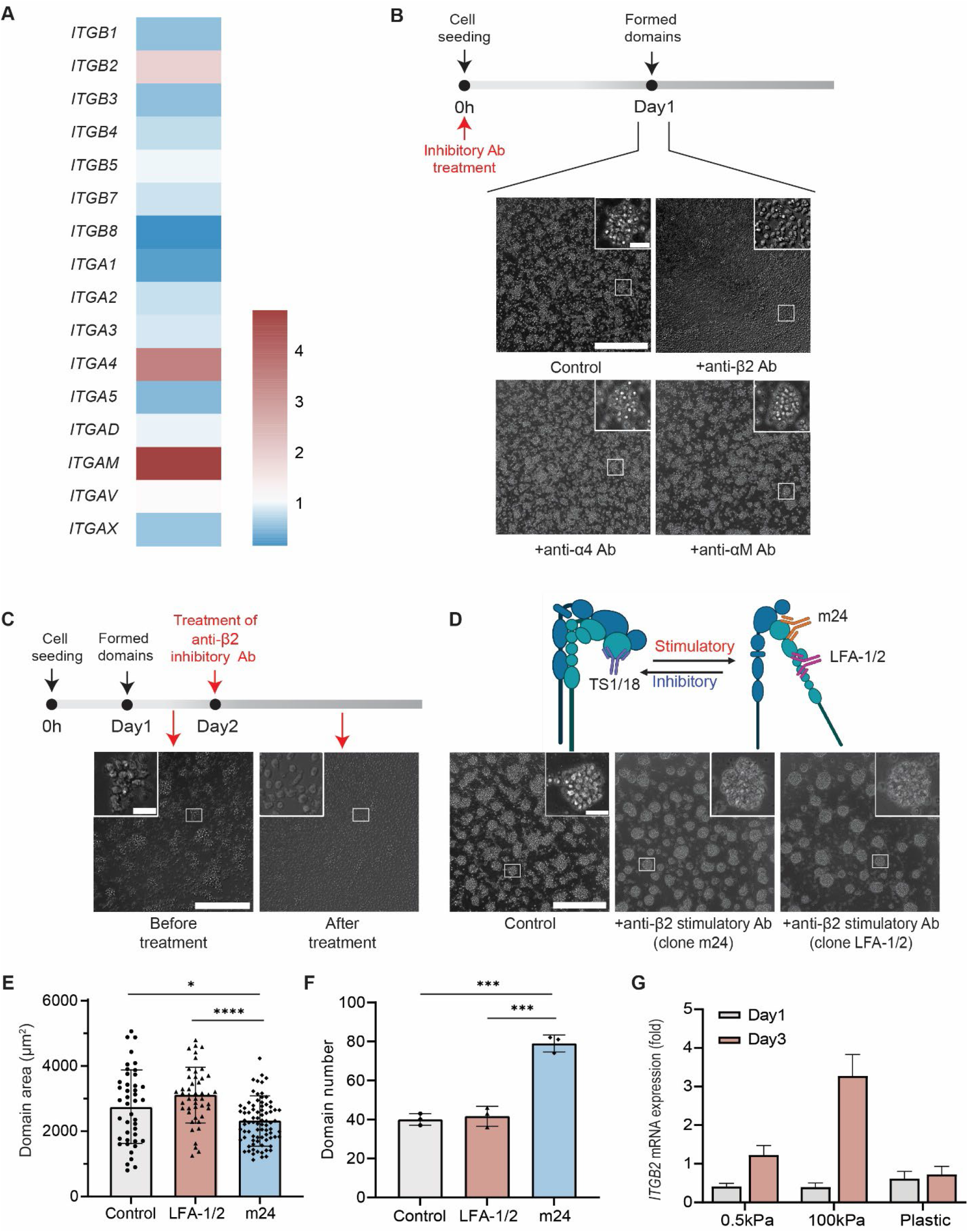
β2 integrin as a mediator of monocyte homotypic domain formation. (**A**) RT-qPCR analysis of cell surface integrins mRNA expression profile using cell samples collected on day1 and day 3 from 0.5 kPa substrate. Relative fold changes of different integrins were measured relative to day 1, including upregulated *ITGB2* (integrin β2), *ITGA4* (integrin α4) and *ITGAM* (integrin αM). (N=3 donors and n=3 wells for a total of 9 experiments per condition). (**B**) Representative images of domain formation when monocytes were treated with inhibitory antibodies against integrins β2, α4 and αM during seeding. (**C**) Representative images of reversible domain collapsing on day 2 after formation when treated with inhibitory anti-β2 antibody. Cartoon showing different binding subunits of inhibitory (clone TS1/18) and stimulatory (clone m24 and LFA-1/2) anti-β2 antibodies. Representative images of induced domain formation when monocytes were treated with stimulatory anti-β2 antibodies. (**E-F**) Characterizations of domain areas and numbers of monocytes when treated with stimulatory (clone m24 and LFA-1/2) anti-β2 antibodies. (**G**) RT-qPCR analysis of *ITGB2* expression trend on three substrates: 0.5 kPa, 100 kPa, and plastic. The mRNA expression trend of *ITGB2* correlated well with the domain development over 3-day period on different substrates. Relative fold changes in expression of different integrins were measured relative to day 0. (N=3, n=3). Phase images were taken using a 20x objective. Scale bars, 300 μm (insets, scale bar, 30 μm). Unpaired student’s t-test was used for statistical analysis. Results are presented as mean ± standard deviation.

To further establish cause and effect, we examined the effect of stimulatory antibodies specific to β2 integrin (clones m24 and LFA-1/2) and found that these activation-specific antibodies significantly accelerated the initiation of monocytes domain formation, resulting in highly organized patterns after overnight culture (Fig. 3D). Interestingly, when stimulatory β2 integrin antibody was added, domain area showed a slight decrease while the number of domains showed a slight increase (Fig. 3, E and F), This suggests a decreased number of non-aggregated (single) monocytes on the substrates. In sum, both inhibitory and stimulatory anti-β2 antibodies demonstrated the direct involvement of β2 integrin in the domain formation of monocytes on soft matrix.

Since monocyte domain formation and progression differed for the tested substrate stiffnesses, we hypothesized that the level of expression of β2 integrin on monocytes placed on different substrates would vary accordingly. RT-qPCR analysis was performed on samples collected on days 1 and 3 (relative fold changes normalized to day 0), representing the initialization stage and stabilization/collapse stage of domains on 0.5 kPa, 100 kPa, and plastic collagen-coated substrates. Despite a slight downregulation of β2 integrin on day 1, elevated mRNA expression of β2 integrin on 0.5 and 100 kPa substrates but not on plastic substrate on day 3 correlated well with the mechano-induced domain pattern maintenance shown in Fig. 1C (Fig. 3G).

### Monocytes secrete global inhibitors

Global phase seperation of the monocyte/buffer system into a large monocyte-rich domain and a monocytes-poor domain did not occur, even for long obervation times (> 3 days). Since steady state domain patterns formed instead of multiple domains merging into a large one, we proposed the existence of monocyte-secreted inhibitory soluble factors that partially inhibited inter-cellular adhesion in the system, referred to “global inhibition”. To ascertain this hypothesis, conditioned medium from homotypically aggregated monocytes were harvested and applied on monocytes freshly seeded on a 0.5 kPa substrate. As depicted in Fig. 4A, monocytes treated with 2x or 4x diluted conditoned medium completely abolished domain formation, presumably due to the high concentration of inhibitory soluble factors secreted by monocytes during the initiation/incubation stage. The monocytes cultured in 8x diluted conditioned medium showed a delayed domain formation and featured significantly smaller domain size and decreased migration speed than control monocytes for overnight incubation (Fig. 4, B and C). As is shown in Supplementary Fig. 2, secreted proteins from aggregated monocytes with a molecular weight around 110-130kDa might be the inhibitory soluble factors that drove the abrogation of domain formation in the presence of condition medium collected from aggregated cells. Identification of the specific inhibitory soluble molecules in the conditioned medium is beyond of the scope of this paper.

**Fig. 4.**
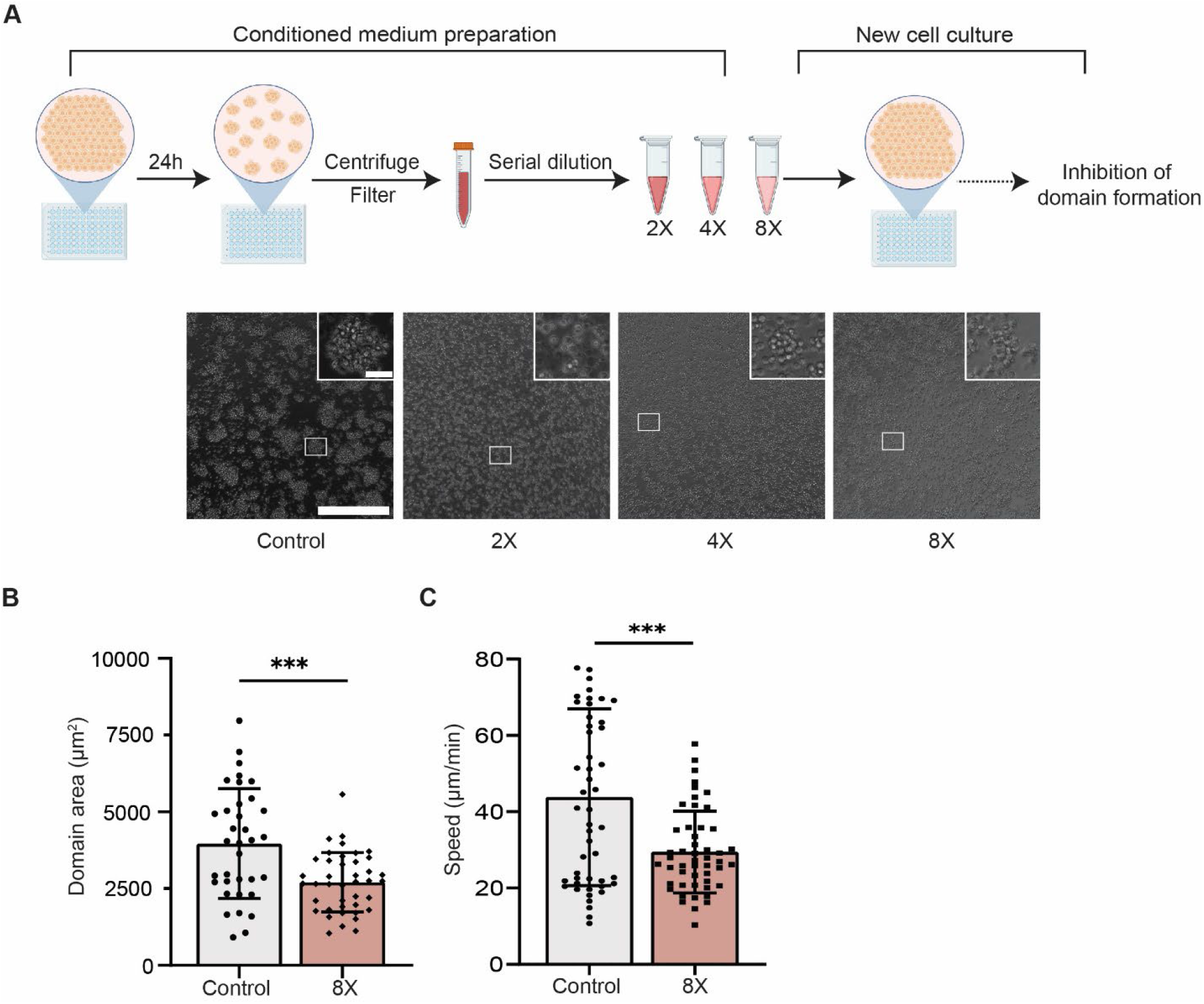
Conditioned media harvested from monocytic domains abrogated the onset of phase separation. (**A**) Conditioned medium harvested from steady state monocyte domains negatively influenced the domain formation initiation in a concentration-dependent manner. Only 8x diluted conditioned medium allowed for monocyte domain formation. (**B-C**) Decreased steady state domain area and cell speed decreased when monocytes were treated with 8x conditioned medium. Unpaired student’s t-test was used for statistical analysis. Results are presented as mean ± standard deviation. Scale bars, 300 μm (insets, scale bar, 30 μm).

### A computational model describes domain formation and predicts key roles for cell migration and seeding density

This spontaneous, reversible, patterned homotypic aggregation of monocytes resembles the motility-induced phase separation (MIPS) of active matter^29, 39, 40^ and displays properties of pattern formation described by the Turing mechanism^41^. To further explore domain formation of monocytes, we developed a corresponding computational model based on the Cahn-Hilliard equation^33, 34^. This model describes the process of phase separation by which the two components (monocytes + surrounding medium) of a binary fluid spontaneously separate and form domains that are pure in each component. Inhibitory factors were introduced to the system following the “global inhibition” theory that we propose and validated. Diffusion, cellular production and degradation of these soluble molecules were combined together to shape their distribution. Considering the relatively consistent initiation time of monocyte domain formation on 0.5 kPa substrate (Supplementary Fig.1), we assumed an elevated affinity of cell adhesion molecules via mechano-sensing as the driving force of phase separation initialization, which we referred to as “local activation”.

The parameters and variables used for modeling the domain formation process are listed in Table 1. Due to the slow proliferation of human primary monocytes, cell growth was ignored when building up the model for simplification (characterization of domain area and number on day 3 of experiments were used to directly compare with steady state results of the simulation). The model consisted of individual cells in an environment that contains diffusible inhibitory molecules. We made the assumption that both cells and inhibitory molecules were diffusible, and we described the free energy of the cells using the equation 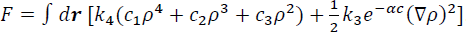 . The first term of this equation is the prototypical bulk free energy of a hard-core particle with short range attraction, which roughly describes cells with mutual adhesion. The second term represents the interfacial energy between cells and the surrounding medium. We hypothesized that the interfacial energy was negatively influenced by inhibitory molecules, following a declining exponential function. Given the free energy, we can obtain the chemical potential as: 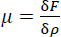. The flux is then given by *j* = −*D*_1_ ∇μ and the final equation for cell movement is described by: 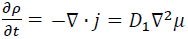. The production and movement of inhibitory factors are described by a reaction-diffusion equation. The final governing equations for both cells and inhibitory factors are:

**Table 1.** Parameters and variables of the computational model.

| Parameter & variable | Dimensionless form | Description | Base value (dimensionless) |
| --- | --- | --- | --- |
| $\rho$ | $\tilde{\rho} = \rho L^2$ | Cell density | / |
| $c$ | $\tilde{c} = c L^2$ | The concentration of inhibitory factor | / |
| $D_1$ | $\tilde{D}_1 = \frac{D_1 T}{L^2}$ | Diffusion coefficient of cells | 1 |
| $D_2$ | $\tilde{D}_2 = \frac{D_2 T}{L^2}$ | Diffusion coefficient of inhibitory factor | 20 |
| $k_1$ | $\tilde{k}_1 = \frac{k_1 T}{L^2}$ | Production coefficient of the inhibitory factor | 1 |
| $k_2$ | $\tilde{k}_2 = k_2 T$ | Degradation coefficient of the inhibitory factor | 0.5 |
| $k_3$ | $\tilde{k}_3 = \frac{k_3}{L^2}$ | Coefficient of interfacial energy | 0.5 |
| $k_4$ | $\tilde{k}_4 = k_4$ | Coefficient of bulk free energy | 2.5 |
| $c_1$ | $\tilde{c}_1 = c_1 / L^4$ | Coefficient in bulk free energy | 0.1024 |
| $c_2$ | $\tilde{c}_2 = c_2/L^2$ | Coefficient in bulk free energy | -0.4551 |
| $c_3$ | $\tilde{c}_3 = c_3$ | Coefficient in bulk free energy | 0.5333 |
| $\alpha$ | $\tilde{\alpha} = \alpha/L^2$ | Coefficient of “global inhibition” | 0.5 |
| $l$ | $\tilde{l} = l/L$ | Side length of the square region | 100 |

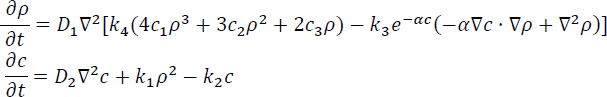

Here, *ρ* and *c* are the cell density and concentration of inhibitory molecules, respectively. We solved the equations in a square region with side length *l*. In our model, we assumed no flux at the boundary for both cells (*ρ*) and inhibitory molecules (*c*). The initial cell density was set with a random small perturbation around a homogeneous state following a uniform distribution. The initial inhibitory molecule concentration was set as zero. To better match the domain area and number characterization results of simulations to experiments conditions, we further defined dimensionless parameters by setting characteristic length (*L*) and time (*T*) scales (Table 1). All the normalized variables and parameters in the equation are defined as: 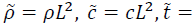 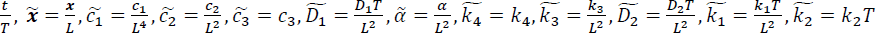. The dimensionless parameters were fixed at base values (Table 1) in all simulations if not specified. The initial cell density was set as *ρ̃* ∼(0.7,0.8) if not specified. In all simulations, the length and time scales are set as: *L* = 8*μm*, *T* = 0.012*h*.

To validate the “global inhibition” theory that we proposed, we first simplified the model by setting parameters *k͠*_1_, *k͠*_2_ as 0 to ignore the production and degradation of the inhibitory diffusible factors. As shown in Fig. 5A-B, with all other parameters fixed, when no inhibitory factors existed in the system (*c̃* = 0), scattered domains eventually merged into larger domains at steady state, which increased the mean domain area while decreased the domain number compared to scenario where inhibitory factors were set at a constant concentration (*c̃* = 2) across the whole simulation. Without inhibitory factors, domain area reached 8543 µm^2^ at steady state, which is significantly larger than the experimental domain area value at day 3 of 4771 µm^2^ (Fig. 1C). Similarly, when tuning up the coefficient of interfacial energy *k͠*_3_ that reflects higher intercellular adhesion activity to simulate the “local activation” (production, degradation of inhibitory factors included), the mean domain area increased to 3809 µm^2^ and decreased the number of domains, which matched observed experimental values (Fig. 5, C and D).

**Fig. 5.**
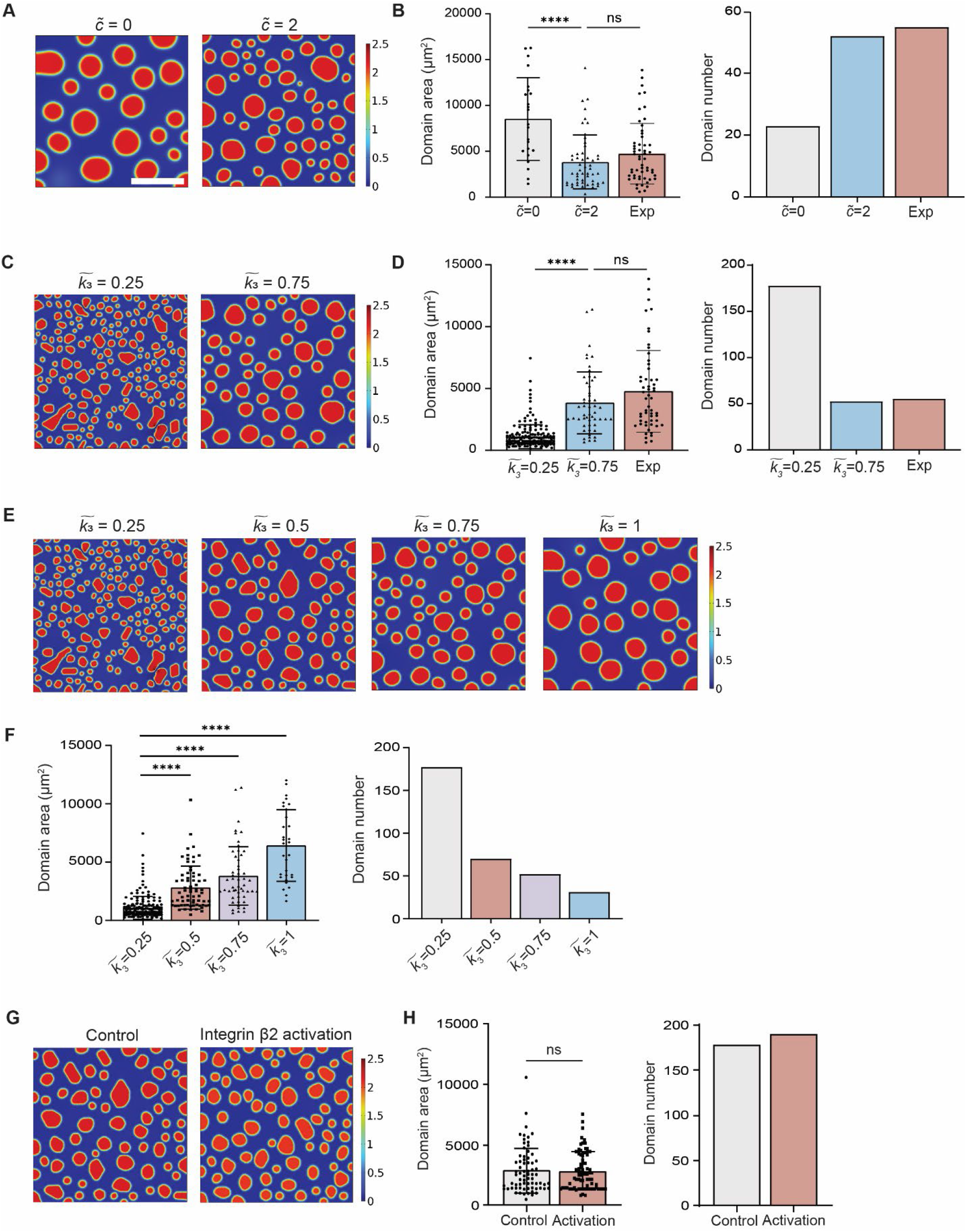
Validation of “global inhibition, local activation” in the simulation model via tuning of single parameters. (**A**) Effect of inhibitory factors on domain formation simulation at constant concentration of *c̃* = 0 and *c̃* = 2 (no production and degradation of inhibitory factors was considered). (**B**) Mean domain area decreased while domain number increased as the concentration of inhibitory factor increased, which matched the experimental characterization results. (**C**) Effect of interfacial energy coefficient on domain formation simulation at *k͠*_3_ = 0.25 and *k͠*_3_ = 0.75. (**D**) Increase in mean domain area and decrease in domain number to similar level of experimental characterization results was observed when higher interfacial energy coefficient was applied. (**E-F**) Increase of interfacial energy coefficient (*k͠*_3_) along with other key parameters fixed resulted in a monotonic increase in domain area and decrease in domain number. (**G-H**) By increasing interfacial energy coefficient (*k͠*_3_) and adjusting the bulk free energy coefficients (*c̃*_1_, *c̃*_2_, *c̃*_3_ to (0.1245, −0.5269, 0.5880), the mean domain area remained unchanged while domain number increases, which is consistent with experiment findings. In the simulation, all parameters except *k͠*_3,_ *c̃*_1_, *c̃*_2_, *c̃*_3_ were fixed. Results were obtained at t = 24h, representing the steady state. Unpaired student’s t-test was used for statistical analysis. Results are presented as mean ± standard deviation. Scale bar = 300 μm.

After identifying β2 integrin as the “local activation” factor in domain formation, we were set to recapitulate the observed β2 integrin stimulation phenomenon (Fig. 3D-F) in our proposed computational model. We first adjusted the interfacial energy coefficient *k͠*_3_ while fixing all other parameters to prove the independence of one key parameter’s influence on the simulation results. As the β2 integrin activity increased, there was a monotonic increase in the domain area and a decrease in the number of domains at steady state (Fig. 5, E and F). Then by increasing the interfacial energy coefficient *k͠*_3_ and adjusting accordingly the bulk free energy coefficients *c̃*_1_, *c̃*_2_, *c̃*_3_ to simulate the experimental setting, a similar trend of unchanged domain area and increased domain number was achieved at steady state (Fig. 5, G and H).

Taken together, we successfully built a Cahn-Hilliard equation-based computational model that closely matched the observed homotypic domain formation.

In addition to inhibitory soluble factor concentration *c̃* and interfacial energy coefficient *k͠*_3_, the cell diffusion coefficient *D͠*_1_ (cell motility) and the cell density *ρ̃* are two key parameters in the computational model that may affect the steady state simulation results. Here we provide predictions of how these two parameters may influence the monocyte domain formation trend. When gradually tuning up the diffusivity coefficient of monocyte *D͠*_1_, the model predicted that the steady state domain area increased while decreasing the number of domains (Fig. 6, A and B). Interestingly, a plateau was reached when *D͠*_1_ became > 20, indicating that locally concentrated inhibitory molecules around formed domains were sufficient to block the merging of adjacent monocyte clusters, no matter how migratory the cells were. In terms of cell density, considering that the coefficients (e.g., interfacial energy coefficient *k͠*_3_) in the free energy may be a function of cell seeding density^33^ and single-cell level integrin β2 expression was positively correlated with cell density (Supplementary Fig. 3), we increased the coefficient of interfacial energy *k͠*_3_ correspondingly while increasing the cell density *ρ̃*. A monotonic increase of *ρ̃* in the simulation resulted in increased domain area (Fig. 6C). Interestingly, when high cell density *ρ̃* was applied, relatively consistent domain numbers were observed at steady state, which indicates that cell density in a single domain remained constant.

**Fig. 6.**
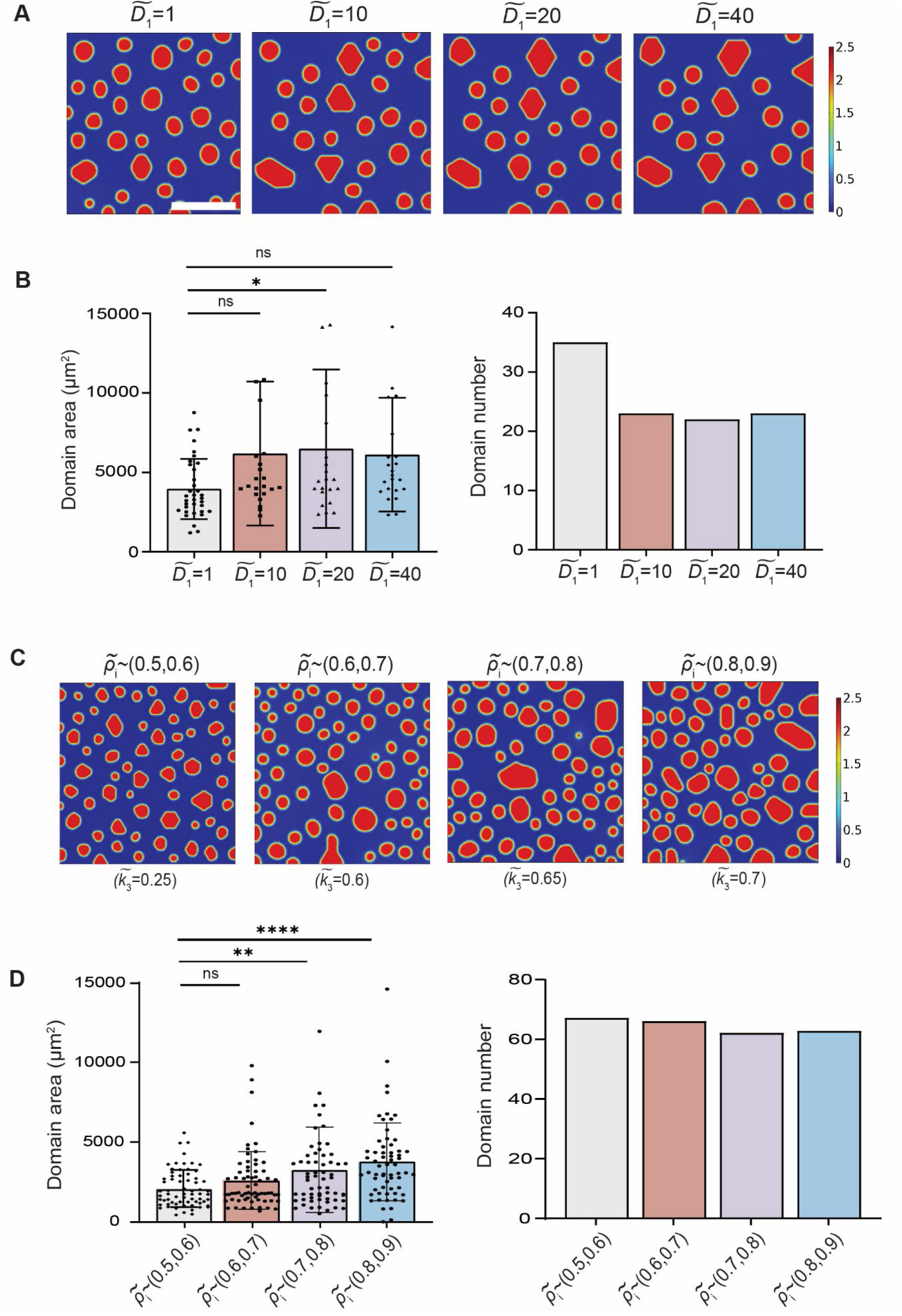
Model prediction on cell motility and seeding density’s effect on monocyte domain formation. (**A**) Simulation results illustrating steady state domain formation with monotonically increased diffusion coefficient *D͠*_1_. All parameters except *D͠*_1_ were fixed. (**B**) Characterization of domain area and domain number at different cell diffusion coefficient *D͠*_1_. Domain area increased with decreasing domain number, which plateaued at *D͠*_1_ = 20. (**C**) Simulation results illustrating steady state domain formation with monotonically increased initial cell density *ρ̃*. Coefficient of interfacial energy *k͠*_3_ were changed correspondingly with different cell density *ρ̃* to reflect the energy density as a function of cell density. (**D**) Characterization of domain area and domain number at different initial cell density *ρ̃*. All parameters except *ρ̃* and coefficient of interfacial energy *k͠*_3_ were fixed. Results were obtained at t=24h, representing the steady state. Unpaired student’s t-test was used for statistical analysis. Results are presented as mean ± standard deviation. Scale bar, 300 μm.

In the following sections, we carried out additional experiments to verify the accuracy of these model predictions.

### Influence of random and chemokine-directed cell motility on domain formation

Just as motility-induced phase separation of active matters^39, 40^, the formation of domains observed in our study was driven by the active movement of monocytes on the substrates. Previous studies have categorized immune cell migration into two distinct modes: chemotaxis and random migration^42^. To understand whether the motility of monocytes could influence the domain formation process and whether the trend matched the model’s prediction, we treated monocytes with different inhibitors targeting either basal migration or chemotaxis pathways.

Five inhibitors that targeted various cell migration pathways were selected to interfere with monocyte basal random migration. These include inhibitors for ROCK (Rho-associated protein kinase), Myosin, Arp2/3, STAT3, and NHE (Na+/H+ ion exchanger)^43–45^.Inhibitions of ROCK, Myosin, and NHE all led to a significant decrease in the average domain area compared to the control group (Fig. 7A). Accordingly, cell motility was negatively influenced by ROCK and NHE inhibition (Fig. 7B). No significant difference was found in terms of domain number (Fig. 7C). Together, these results indicate that domain size is correlated with cell motility. Slower cell migration results in smaller domain size. Consistent with the random migration inhibitor experiments, decreased diffusion coefficient *D͠*_1_ resulted in inadequate domain development in simulation with more domains with smaller area (Fig. 7, D and E).

**Fig. 7.**
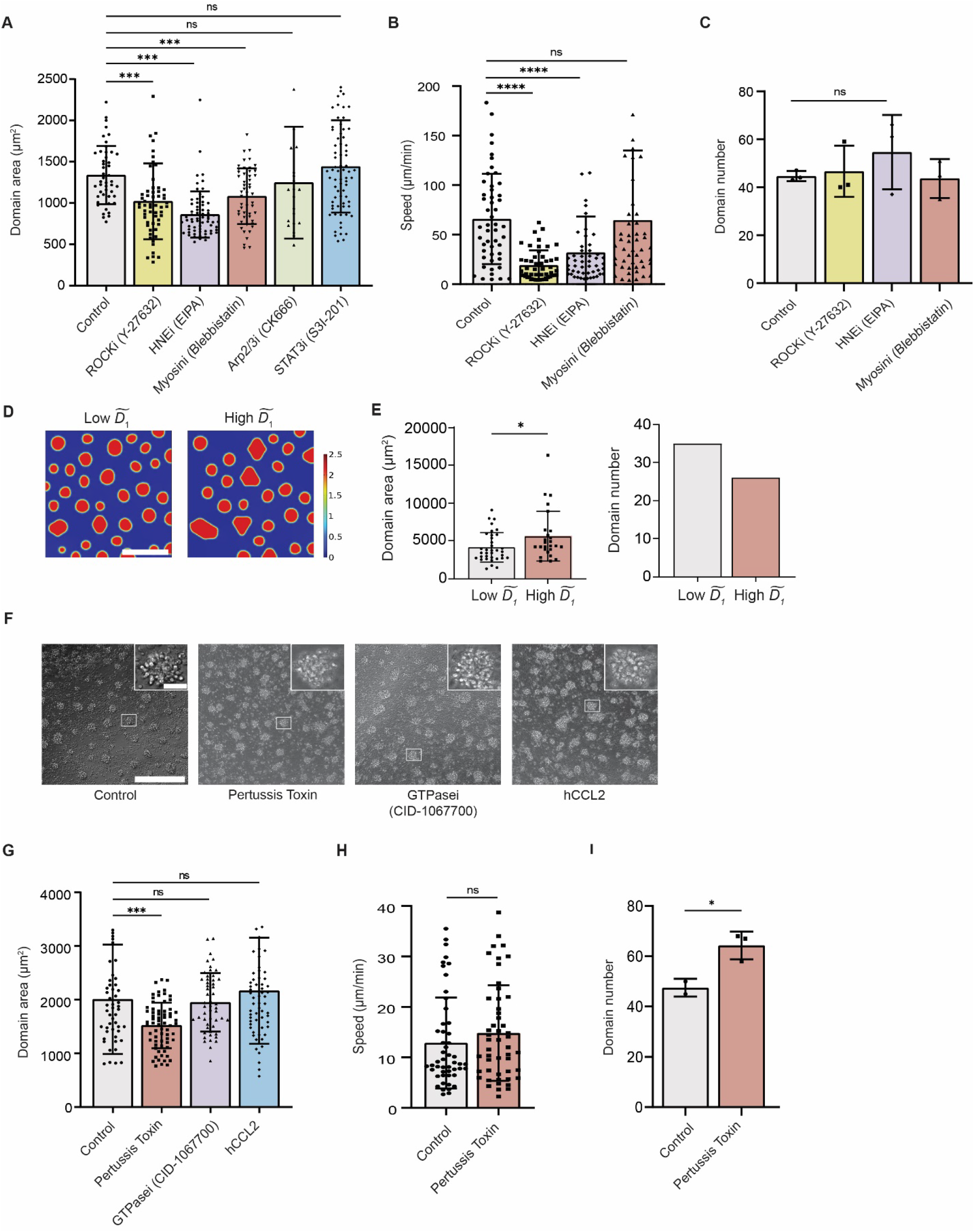
Modulation of monocyte homotypic domain formation by cell motility. (**A**) Monocytes seeded on 0.5 kPa substrate were treated with 20 μM ROCK inhibitor Y-27632, 100 μM Arp inhibitor CK666, 100 μM STAT3 inhibitor S3I-201, 10 μM Blebbistatin, and 10 μM EIPA. The average domain area significantly decreased following Y-27632, blebbistatin, and EIPA treatments. (**B**) Y27632 and EIPA treated monocytes showed significantly reduced cell motility. (**C**) No significant differences existed in domain numbers of Y27632, EIPA and Blebbistation treated conditions compared to control. Increased domain number when treated with EIPA was observed, though. (**D**) Representative simulation results comparing cells with low and high motility. In the simulation, all parameters were fixed except *D͠*_1_. (**E**) Domain area decreased in the low cell motility group, while low cell motility led to an increase in domain number. (**F**) 20x representative images on monocyte domain formation when treated with complete medium, pertusis toxin, GTPasei and hCCL2, respectively. (**G**) Evaluation of domain area in the presence of three molecules targeting monocyte chemotaxis. Monocytes on 0.5 kPa substrate treated with pertussis toxin formed significantly smaller domains compared to the control group. (**H**) Cell motility did not significantly decrease upon pertussis toxin treatment. (**I**) Pertussis toxin treated monocytes showed increased domain number. Unpaired student’s t-test was used for statistical analysis. Results are presented as mean ± standard deviation. Scale bar = 300 μm for both experimental phase images and simulations (insets, scale bar, 30 μm).

To further validate the computational model in which diffusion of monocytes is considered as the sole driving force for domain formation instead of chemotaxis, we employed pertussis toxin (general chemotaxis inhibitor), CID-1067700 (a pan-GTPase inhibitor) and hCCL2 (block CCL2-CCR2 axis) to target monocyte chemotaxis. For these three treatments, monocytes were only found to aggregate into significantly smaller size (1522 ± 425 μm^2^) under the influence of pertussis toxin compared to the non-treated control group (2007 ± 1020μm^2^) (Fig. 7, F and G), which also resulted in higher domain number (Fig. 7I). However, pertussis toxin treatment did not significantly reduce cell movement compared to the control groups (Fig. 7H). These findings suggest that chemotaxis may not be the driving force behind monocyte aggregation in the domain formation process and diffusion along is sufficient to describe monocyte motion in the model.

### Onset of domain formation is independent of cell seeding density

To eliminate the possibility that monocyte domain formation is caused by excessive cells in each well, we adjusted the initial cell seeding density in each well. Three different seeding densities – 50,000 cells/well, 25,000 cells/well, and 10,000 cells/well – were used, which we referred to as high, medium, and low densities. On day 1, we observed that monocytes tended to form larger domains at a high cell density. The high-density group resulted in larger domains (2396 ± 996 μm^2^) compared with the low-density group (992 ± 423 μm^2^) (Fig. 8A). A linear correlation between the average monocyte domain area and the cell seeding density was observed (Fig. 8B). However, despite the variations in initial cell density, we consistently observed the initiation of domain formation across all three density groups (Supplementary Video 5). This suggests that the process of domain formation is independent of the initial monocyte seeding density, as evidenced by the formation of domains even in the low-density group where cells were initially spaced far apart.

**Fig. 8.**
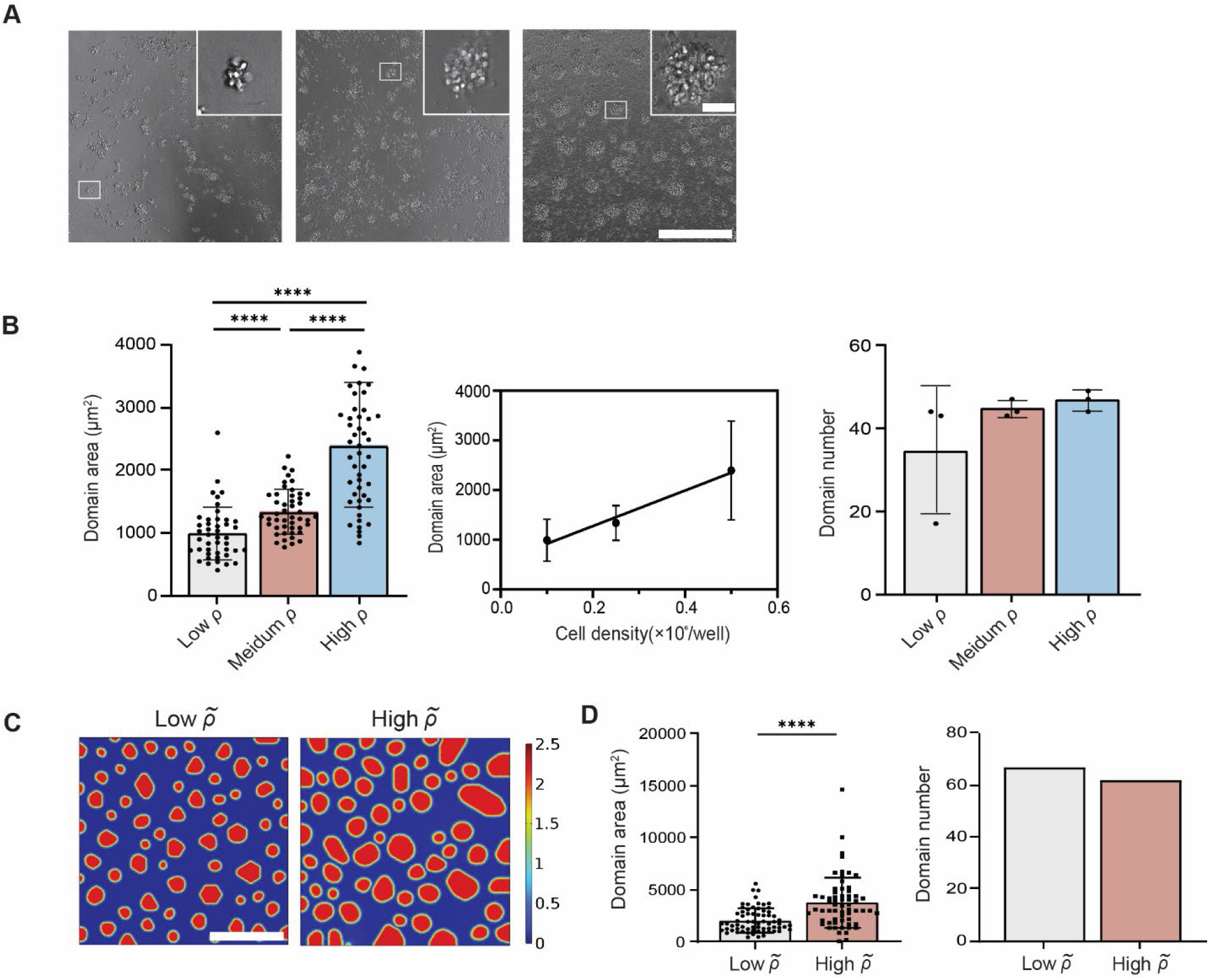
Monocyte homotypic domain formation was modulated by local seeding density. Three cell seeding densities (50,000, 25,000, 10,000 cells/well) representing high, medium, and low density were used. (**A**) Representative 20x images of three cell seeding density captured on day 1. (**B**) The averaged domain area on day 1 exhibited a linear correlation with cell seeding density. (**C**) Representative simulation results of low- and high-density groups (*ρ̃* ∼ *U* (0.5,0.6) and *ρ̃* ∼ *U* (0.8,0.9)). All parameters except cell density *ρ̃* were fixed. (**D**) Significant increase of domain area was observed when comparing high seeding density group condition to low seeding density condition, while the high-density group exhibited a slight decrease in domain number. In the simulation, all parameters except initial cell density were held consistent. Results were obtained at t=24h, representing the steady state. Unpaired student’s t-test was used for statistical analysis. Results are presented as mean ± standard deviation. Scale bar = 300 μm for both experimental phase images and simulations (insets, scale bar = 30 μm).

Bumping up the cell density from *ρ̃* ∼(0.5,0.6) to *ρ̃* ∼(0.8,0.9) in our computational model to mimic the low and high cell densities (25,000 *vs*. 50,000 cells/well) showed a significant increase in domain area (Fig. 8, C and D). Decreased domain number was observed, but the deviation wasn’t large. In general, the simulation results matched up well with the experimental outcomes with few concerns that remain to be solved. For example, although the initial diffusible factors concentration *c̃* was set at 0, higher cell density *ρ̃* introduced considerable amount of inhibitory soluble factors that might interfere with the steady state simulation results. A time-delayed introduction of inhibitory soluble factors may offer a better solution for future optimization of the model.

## Discussion

Monocytes originating from bone marrow are released into the bloodstream and recruited to tissues to fight against pathogens upon maturation. In tumors, monocytes can aggregate into dynamic complex structures (“hot spots”) *in vivo* (e.g., Fig. 1A). Here, we demonstrate that monocytes respond to differences in matrix stiffness. On a matrix of (low) physiological stiffness, the ensuing activation of β2 integrin on the monocyte surface, which initiates intercellular adhesion, and the secretion of global inhibitors together produce local cellular phase separation, resulting in patterned domain formation and maintenance.

The observed homotypic immune cell aggregates are reminiscent of dynamic phases observed in active matter systems. In particular, flocking transitions seen during collective movement of fish and birds also produce high density aggregates^27, 28^. More recent theoretical work has revealed rich phase behavior of self-propelled particles in terms of the underlying microscopic interactions^29, 30^. Different from flocking, which are governed by local short-range interactions between self-driven particles, cells possess both short range interaction and long-range signals, and therefore potentially can generate new complex behavior. Cells can be mechanosensitive and secrete factors. Moreover, cell movement is also distinctly different from propelled particles, and are instead described by persistent random walks (PRW)^31, 32^.

Our functional studies revealed a reason for monocytes to aggregate on matrices of physiological stiffness: enhanced survival accompanied with decreased phagocytic capability. This optimization scheme is unique to living cells, and fundamentally different from active matter considered thus far^39, 40^. Moreover, self-propelled colloidal particles neither secrete inhibitory molecules nor actively modulate adhesion molecules on their surface, which are two key ingredients leading to our observed patterned aggregates. Still, the resemblance between this spontaneous, reversible, patterned aggregation of monocytes and the motility-induced phase separation (MIPS) of active matters motivated us to build a corresponding computational model based on the Cahn-Hilliard equation. Secreted inhibitory soluble molecules, β2 integrin and cell motility were identified as key ingredients of the model, and all were validated by matching simulations with experimental results.

Previous studies on monocytic cell line U937 reported the involvement of LFA-1/VLA-4 in their homotypic adhesion on tissue culture plastic. Addition of stimulatory LFA-1/VLA-4 antibodies was found to induce the initiation of cell aggregation^46, 47^. However, the standard *in vitro* culture methods using tissue-culture plastics hardly provide physiological-relevant mechanical properties, which can dramatically shape the phenotype and functionality of monocytes, as shown in this work. Biological implications of this work remain to be tested *in vivo*, but the biophysical mechanism presented in this paper suggests that, in the case of tumor onset and progression, hot spots formation is initiated in inflamed (soft) tissues (stiffness, 1kPa) containing precursor lesions and immune cells, not after cancer cells and cancer-associated fibroblasts deposit and crosslink collagen and other extracellular matrix molecules^48^, rendering the tumor matrix much stiffer (>25 kPa)^35^.

Future iterations of the model are required to improve the accuracy of simulation against the experimental results. For example, in contrast with our current simplification that cells produce inhibitory soluble factors at a constant rate from the beginning, time-delayed production of inhibitory factors based on the separation level (e.g., the characteristic distance between adjacent domains) can be applied so that high cell density won’t introduce extremely high level of inhibitory soluble factors that might interfere with the normal initialization of monocyte domain formation. In addition, based on an active colloid-like system, the cellular complexity and heterogeneity was not fully taken into consideration. Our model does not take into account that single-cell level β2 integrin expression is positively correlated with cell seeding density. Similarly, aggregated monocytes may exhibit a different secretomic profile compared to actively migrating cells and potential effects of layer-stacking domain structure on characteristic domain areas are yet left unsolved in the current iteration of our model, which urges the incorporation of equations describing related cellular functions.

## Acknowledgments

The authors would like to thank all members of the Wirtz lab and Sun lab for their feedback. This work was partially supported through grants to D.W. from the National Cancer Institute (U54CA143868 and U54CA268083), the National Institute of Arthritis and Musculoskeletal and Skin Diseases (U54AR081774), and the National Institute on Aging (U01AG060903).

## Author contributions

W.D., J.Z. designed and carried out the experiments. Y. W. developed the simulation model and carried out all simulations. The manuscript was written by W. D., Y.W. with edits and input from D.W., S.S. and J.Z.

## Methods

### Isolation and culture of human primary monocytes / macrophages / B cells / NK cells / T cells

PBMCs were isolated from donors’ whole blood by Ficoll-paque PLUS (cytiva) density gradient centrifugation. Human primary classical monocytes (CD14^+^CD16^-^) were further isolated using the Classical Monocyte Isolation Kit (Miltenyi Biotec) by magnetic-activated cell sorting (MACS). Human CD14^+^CD16^-^ monocytes were resuspended in DMEM (Corning) supplemented with 4.5g/L glucose, 10% v/v heat-inactivated fetal bovine serum (Corning), and 1% v/v penicillin-streptomycin (Sigma). Monocytes were cultured at a density of 0.5×10^6^ cells per well and incubated at 37 °C. 96-well culture plates, either standard tissue culture plastic coated with collagen I or Softwell 96-well plates (Matrigen) containing collagen I coated polyacrylamide gel with stiffness values of 0.5 and 100 kPa were used. Similarly, human primary B cells / NK cells / T cells were isolated from donor PBMCs using corresponding isolation kits (B cell -- STEMCELL 17954; NK cell -- STEMCELL 17955; T cell -- STEMCELL 17911) per manufacturer protocol. B cells were cultured in ImmunoCult-XF B cell base medium (STEMCELL) with B cell expansion supplement (STEMCELL). NK cells were cultured in ImmunoCult NK cell base medium (STEMCELL) with NK cell expansion supplement (STEMCELL). T cells were cultured in X-VIVO 15 medium (Lonza) with 10% v/v FBS (Corning), IL-2 (R&D) and CD3/CD28 T cell activator (STEMCELL). Macrophages were differentiated from negatively isolated human primary monocytes in macrophage attachment medium -- DMEM (Corning) supplemented with 4.5g/L glucose, 10% v/v heat-inactivated fetal bovine serum (Corning), 1% v/v penicillin-streptomycin (Sigma) and 50 ng/mL human recombinant M-CSF (R&D). Monocytes were seeded at a density of 2.5×10^6^ in each well of a standard 6-well tissue-culture plate (5×10^6^ monocytes per well) in macrophage attachment medium the day they were isolated from donor PBMCs. After 3 days, cells remaining in suspension were gently aspirated away and fresh macrophage attachment medium were added. Macrophages were ready to be harvested and seeded upon 7 days of differentiation, when they were regarded as M0 naïve macrophages.

### Collagen I coating of tissue-culture plates

Based on the previously described method, low-concentration Collagen I (Corning 354249) was diluted with 0.1% acetic acid to a final concentration of 100 μg/mL. 96-well plastic cell culture plates were washed with PBS 3 times before adding 32 μL diluted collagen I per well. After 1 hour of incubation, excess liquid in each well was aspirated out carefully followed by PBS washing. Complete DMEM medium was used to condition the plate for 1 hour before use.

### Antibody inhibition of monocyte cell surface integrins

To explore the cell surface molecule engaged in the domain formation process, 6.25 μg/mL blocking antibodies against Integrin β2 (clone TS1/18, Biolegend) and 6.25 μg/mL activation antibodies against Integrin β2 (clone m24 and LFA1/2, Biolegend) were added in cell culture medium at the start of the culture.

### Chemical Inhibition of monocyte migration

Isolated human CD14^+^CD16^-^ monocytes were incubated with various inhibitors upon seeding. Several inhibitors were used to interfere with the random migration of monocytes: 20 μM RhoA Kinase ROCK inhibitor Y-27632 (Selleckchem), 100 μM Arp inhibitor CK666 (Selleckchem), 100 μM STAT3 inhibitor S3I-201 (Selleckchem), 10 μM Blebbistatin for non-muscle myosin II inhibition (Millipore Sigma), and 10 μM EIPA (Millipore Sigma) to block the sodium-hydrogen exchanger (NHE). Monocytes were incubated with 2μg/mL Pertussis toxin (EMD Millipore Crop), 20 μM CID-1067700 (MedChemExpress), and 2 μg/mL recombinant human CCL2 (Biolegend).

### PrestoBlue cell viability assay

2x working solution of PrestoBlue was prepared by mixing 10x stock solution to complete DMEM medium at a ratio of 1:5. 100µL of medium was gently aspirated from each testing well of a 96-well plate (200µL in total) and 100µL of 2x working solution was added slowly to avoid any agitation to the monocyte domains. After 3 hours of incubation in a 37 ° C, 5% CO_2_ incubator, the plate was wrapped in aluminum foil and transferred to SpectraMax M3 plate reader (Molecular Devices). Fluorescence signals were read at an excitation of 560nm and emission of 590nm.

### Flow cytometry

Cell samples were washed 3 times in PBS and resuspended at a concentration of 1 million per mL. Cell suspensions were blocked with Human TruStain FcX (Biolegend) for 15 min under room temperature. The antibody staining solution was then added and incubated at 4 °C for 30 min. Antibodies used for cell labeling were as follows: APC anti-human CD18 (clone LFA-1/2, Biolegend), and Propidium Iodide solution (Biolegend). Wash cells and then resuspend cells in 350 μL FAC wash buffer (1X DPBS containing 5% FBS, 1mM EDTA). Immunofluorescence-stained cells were analyzed on a FACS Canto. Analysis was performed with Flowjo Software version 10.4. The cell surface marker mean fluorescence intensity of each sample was corrected with the mean fluorescence intensity measured for corresponding isotype control.

### Gene expression analysis using RT-qPCR

Total RNA from monocytes subjected to different stiffness conditions and culture time was isolated using RNeasy Micro Kit (QIAGEN). cDNA synthesis was performed using iScript cDNA Synthesis Kit (Bio-Rad). Real-time PCR reactions were set up using iTaq Universal SYBR Green Supermix (Bio-Rad) and were executed in a thermal cycler (CFX384^TM^ Real-Time System, Bio-Rad). The primers designed for specific gene amplification are listed in Table 2. Relative quantitation was performed using the △△Ct method in CFX Manager software.

**Table 2.**
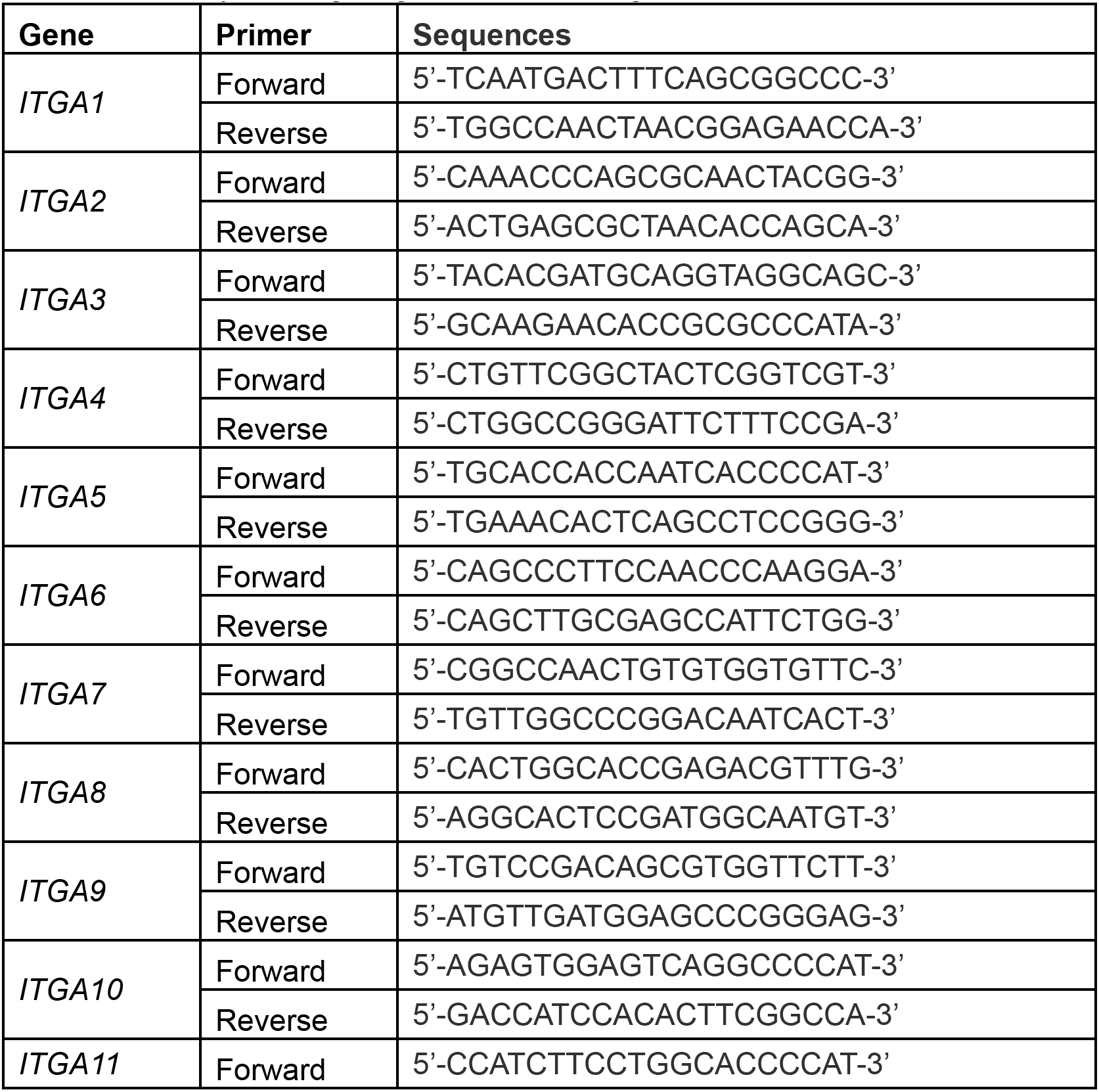
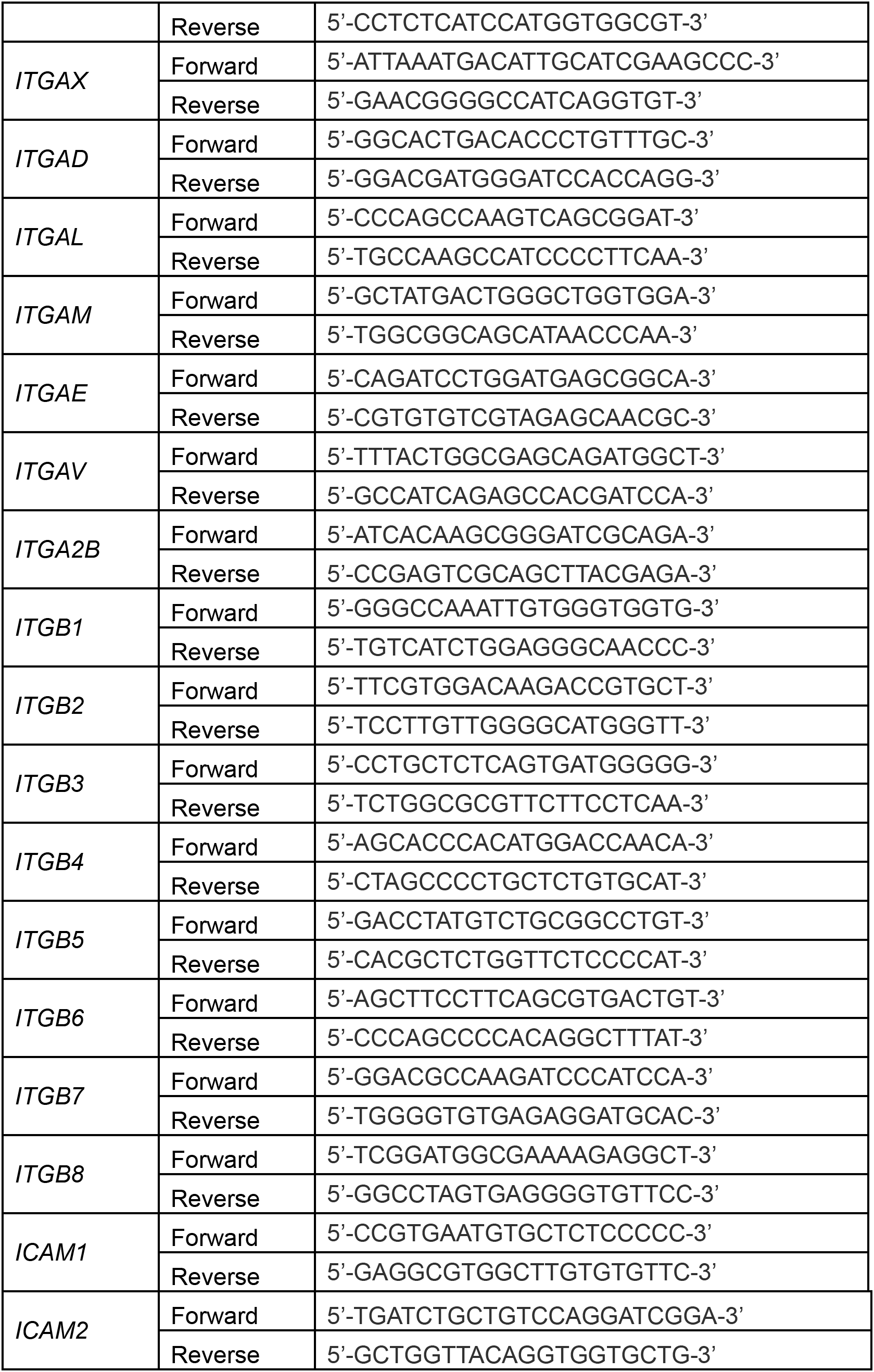
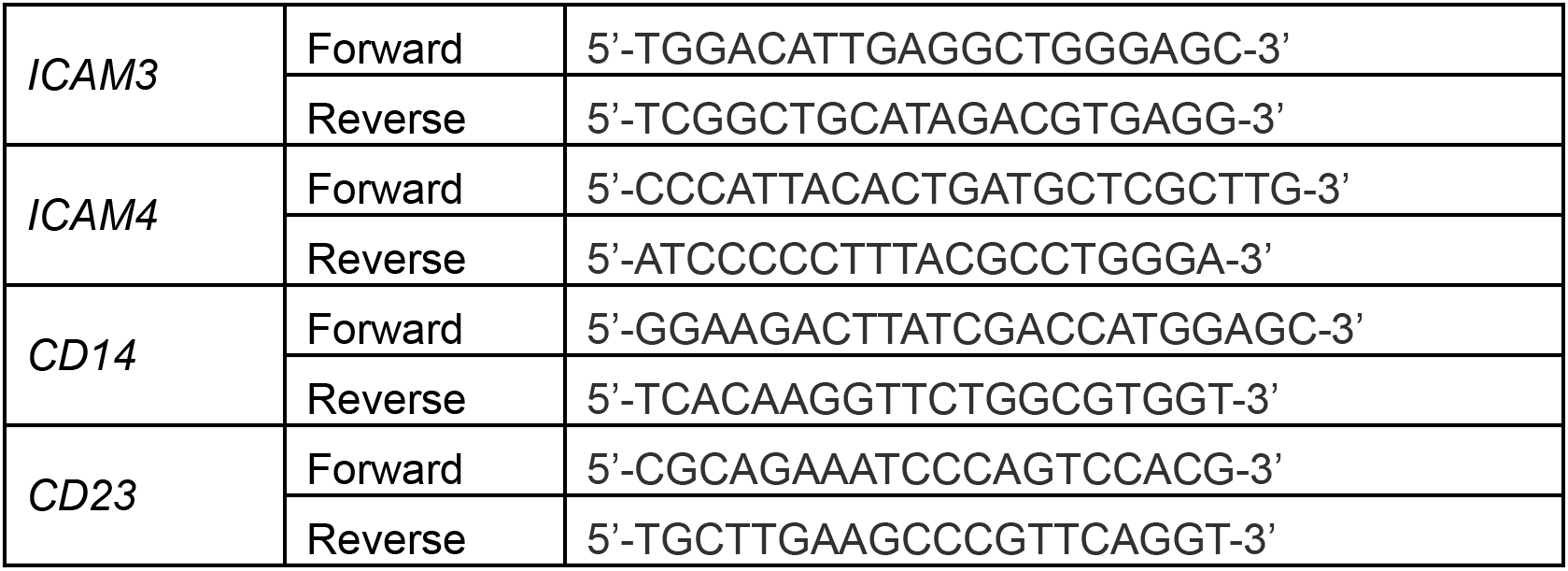
Primer pairs targeting cell surface integrins.

### Western Blot

Whole-cell protein lysates were prepared in clear sample buffer (0.5M Tris pH 6.8, 20% SDS, 50% glycerol in water). Total protein concentration was evaluated using Micro BCA^TM^ Protein Assay Kit (Thermo Scientific). Based on the calculation, the same amount of protein was loaded on 4-15 % SDS-PAGE gels (Bio-Rad) and ran at a voltage of 180V under room temperature for electrophoresis. Protein bands were transferred to the PVDF membrane using the Trans-Blot Turbo system (Bio-Rad). Membranes were blocked in 5% dry milk in TBST for 60 min at room temperature and incubated with diluted primary antibodies. Membranes were incubated with secondary antibodies for 60 min at room temperature. Then the membrane was analyzed under the ChemiDoc^TM^ XRS+ imaging system (Bio-Rad). Images were analyzed using Image Lab software. The intensity of the targeted protein band was normalized using housekeeping protein. Primary and secondary antibodies were used as follows: GAPDH (14C10) Rabbit mAb, Integrin β-2 (D4N5Z) Rabbit mAb, Anti-Rabbit IgG HRP-linked Antibody. All the antibodies were purchased from Cell Signaling Technology.

### Conditioned medium preparation and treatment

A total of 50,000 cells were seeded into each well of the 0.5kPa 96-well plate. Conditioned media were harvested from each well after 24h or 72h incubation. Harvested supernatant was centrifuged at 1000 rpm for 5min and passed through a 0.22μm filter to eliminate any possible cells in it. Aliquots of conditioned media were stored in the -80 °C freezer until use. Different dilutions of conditioned media were prepared by mixing the conditioned media with fresh DMEM in different volume ratios for further monocyte culture on 0.5kPa substrate.

### Ultra centrifugation and silver staining of conditioned medium

Conditioned media were harvested from monocytes seeded on different substrates at different timepoints and filtered through 0.22µm PES filters (Genesee Scientific). To normalize the loading amounts of total proteins, Western blot on Tryp-LE detached cell were carried out on house-keeping protein GAPDH (14C10) to determine the relative cell numbers for different conditions (cell numbers were assumed to be positively correlated with GAPDH band intensity). Conditioned media were concentrated at the ratio of 30x using Amicon Ultra centrifugal filter units with 10k MWCO (Millipore Sigma). Concentrated conditioned media normalized to cell numbers were loaded on 4-15 % SDS-PAGE gels (Bio-Rad) and ran at a voltage of 180V under room temperature for electrophoresis. After breaking SDS-PAGE gels out from the cassette, silver staining was then carried out per manufacturer’s protocol using Pierce Silver Stain kit (Thermo Scientific).

### Live-cell staining and phagocytosis assay

Monocytes were labeled using Cell Tracker Red (CMTPX, Invitrogen) immediately before seeding. After days of incubation, cells inside and outside the domains were separated by gentle pipetting. These two parts of cells were seeded to a new plastic plate at a density of 10,000 cells/well. Carboxylate-modified polystyrene fluorescent beads (Sigma), 1μm in diameter, were added to the cells at the concentration of 5 beads/cell. Fluorescence images were taken every 30 min for 6 hrs. The merged images were obtained to count the total bead number per cell.

### Live-cell imaging and cell tracking

Human Classical monocytes (CD14^+^CD16^-^) were seeded at 50,000 cells/well on a 96-well plate. Images were taken every 1 min for 6 h using Nikon Eclipse Ti2 equipped with a stage top incubator. Cell movements were analyzed using MetaMorph and MATLAB. To ensure the accuracy of manual tracking, >50 cells in each field of view were analyzed.

### Statistical analysis

Graphpad Prism 9 software was used for statistical analysis. An unpaired two-tailed student’s t-test was performed to evaluate the statistical significance between the two groups. For column analysis of multiple conditions, ordinary one-way ANOVA was performed. Significant values were given in grades P < 0.05(*), P < 0.01(**), P < 0.001(***), P < 0.0001(****).

## References

1 Handorf, A. M., Zhou, Y., Halanski, M. A. & Li, W.-J. Tissue Stiffness Dictates Development, Homeostasis, and Disease Progression. Organogenesis 11, 1–15 (2015). https://doi.org:10.1080/15476278.2015.1019687

2 Guimarães, C. F., Gasperini, L., Marques, A. P. & Reis, R. L. The stiffness of living tissues and its implications for tissue engineering. Nature Reviews Materials 5, 351–370 (2020). https://doi.org:10.1038/s41578-019-0169-1

3 Selman, M. & Pardo, A. Fibroageing: An ageing pathological feature driven by dysregulated extracellular matrix-cell mechanobiology. Ageing Research Reviews 70, 101393 (2021). https://doi.org/10.1016/j.arr.2021.101393

4 Sicard, D. et al. Aging and anatomical variations in lung tissue stiffness. American Journal of Physiology-Lung Cellular and Molecular Physiology 314, L946–L955 (2018). https://doi.org:10.1152/ajplung.00415.2017

5 Phillip, J. M., Aifuwa, I., Walston, J. & Wirtz, D. The Mechanobiology of Aging. Annual Review of Biomedical Engineering 17, 113–141 (2015). https://doi.org:10.1146/annurev-bioeng-071114-040829

6 Chen, H., Cai, Y., Chen, Q. & Li, Z. Multiscale modeling of solid stress and tumor cell invasion in response to dynamic mechanical microenvironment. Biomech Model Mechanobiol 19, 577–590 (2020). https://doi.org:10.1007/s10237-019-01231-4

7 Deng, B. et al. Biological role of matrix stiffness in tumor growth and treatment. Journal of Translational Medicine 20, 540 (2022). https://doi.org:10.1186/s12967-022-03768-y

8 Ishihara, S. & Haga, H. Matrix Stiffness Contributes to Cancer Progression by Regulating Transcription Factors. Cancers (Basel) 14 (2022). https://doi.org:10.3390/cancers14041049

9 Wirtz, D., Konstantopoulos, K. & Searson, P. C. The physics of cancer: the role of physical interactions and mechanical forces in metastasis. Nature Reviews Cancer 11, 512–522 (2011). https://doi.org:10.1038/nrc3080

10 Nakasaki, M. et al. The matrix protein Fibulin-5 is at the interface of tissue stiffness and inflammation in fibrosis. Nature Communications 6, 8574 (2015). https://doi.org:10.1038/ncomms9574

11 Boyette, L. B. et al. Phenotype, function, and differentiation potential of human monocyte subsets. PLOS ONE 12, e0176460 (2017). https://doi.org:10.1371/journal.pone.0176460

12 Ginhoux, F. & Jung, S. Monocytes and macrophages: developmental pathways and tissue homeostasis. Nature Reviews Immunology 14, 392–404 (2014). https://doi.org:10.1038/nri3671

13 Wynn, T. A., Chawla, A. & Pollard, J. W. Macrophage biology in development, homeostasis and disease. Nature 496, 445–455 (2013). https://doi.org:10.1038/nature12034

14 Qian, B. Z. et al. CCL2 recruits inflammatory monocytes to facilitate breast-tumour metastasis. Nature 475, 222–225 (2011). https://doi.org:10.1038/nature10138

15 Kiemen, A. L. et al. CODA: quantitative 3D reconstruction of large tissues at cellular resolution. Nature Methods 19, 1490–1499 (2022). https://doi.org:10.1038/s41592-022-01650-9

16 de Groot, A. E. et al. Characterization of tumor-associated macrophages in prostate cancer transgenic mouse models. The Prostate 81, 629–647 (2021). https://doi.org/10.1002/pros.24139

17 Vijayashankar, D. P. & Vaidya, T. Homotypic aggregates contribute to heterogeneity in B cell fates due to an intrinsic gradient of stimulant exposure. Experimental Cell Research 405, 112650 (2021). https://doi.org/10.1016/j.yexcr.2021.112650

18 Malet-Engra, G. et al. Collective Cell Motility Promotes Chemotactic Prowess and Resistance to Chemorepulsion. Current Biology 25, 242–250 (2015). https://doi.org/10.1016/j.cub.2014.11.030

19 Ng, L. G. et al. Visualizing the neutrophil response to sterile tissue injury in mouse dermis reveals a three-phase cascade of events. J Invest Dermatol 131, 2058–2068 (2011). https://doi.org:10.1038/jid.2011.179

20 O’Flaherty, J. T., Kreutzer, D. L. & Ward, P. A. Neutrophil Aggregation and Swelling Induced by Chemotactic Agents1. The Journal of Immunology 119, 232–239 (1977). https://doi.org:10.4049/jimmunol.119.1.232

21 Mangaonkar, A. A. et al. Bone marrow dendritic cell aggregates associate with systemic immune dysregulation in chronic myelomonocytic leukemia. Blood Advances 4, 5425–5430 (2020). https://doi.org:10.1182/bloodadvances.2020002415

22 Lefkowitch, J. H., Haythe, J. H. & Regent, N. Kupffer cell aggregation and perivenular distribution in steatohepatitis. Mod Pathol 15, 699–704 (2002). https://doi.org:10.1097/01.Mp.0000019579.30842.96

23 Harjunpää, H., Llort Asens, M., Guenther, C. & Fagerholm, S. C. Cell Adhesion Molecules and Their Roles and Regulation in the Immune and Tumor Microenvironment. Front Immunol 10, 1078 (2019). https://doi.org:10.3389/fimmu.2019.01078

24 Schittenhelm, L., Hilkens, C. M. & Morrison, V. L. β2 Integrins As Regulators of Dendritic Cell, Monocyte, and Macrophage Function. Frontiers in Immunology 8 (2017). https://doi.org:10.3389/fimmu.2017.01866

25 Herter, J. & Zarbock, A. Integrin Regulation during Leukocyte Recruitment. The Journal of Immunology 190, 4451–4457 (2013). https://doi.org:10.4049/jimmunol.1203179

26 Imhof, B. A. & Aurrand-Lions, M. Adhesion mechanisms regulating the migration of monocytes. Nature Reviews Immunology 4, 432–444 (2004). https://doi.org:10.1038/nri1375

27 Vicsek, T., Czirók, A., Ben-Jacob, E., Cohen, I. & Shochet, O. Novel Type of Phase Transition in a System of Self-Driven Particles. Physical Review Letters 75, 1226–1229 (1995). https://doi.org:10.1103/PhysRevLett.75.1226

28 Tu, Y. Phases and phase transitions in flocking systems. Physica A: Statistical Mechanics and its Applications 281, 30–40 (2000). https://doi.org/10.1016/S0378-4371(00)00017-0

29 Marchetti, M. C. et al. Hydrodynamics of soft active matter. Reviews of Modern Physics 85, 1143–1189 (2013). https://doi.org:10.1103/RevModPhys.85.1143

30 Takatori, S. C. & Brady, J. F. Towards a thermodynamics of active matter. Physical Review E 91, 032117 (2015). https://doi.org:10.1103/PhysRevE.91.032117

31 Wu, P.-H., Giri, A., Sun, S. X. & Wirtz, D. Three-dimensional cell migration does not follow a random walk. Proceedings of the National Academy of Sciences 111, 3949–3954 (2014). https://doi.org.doi.10.1073/pnas.1318967111

32 Li, B. & Sun, S. X. Coherent motions in confluent cell monolayer sheets. Biophys J 107, 1532–1541 (2014). https://doi.org:10.1016/j.bpj.2014.08.006

33 Cahn, J. W. & Hilliard, J. E. Free Energy of a Nonuniform System. I. Interfacial Free Energy. The Journal of Chemical Physics 28, 258–267 (2004). https://doi.org:10.1063/1.1744102

34 Lee, D. et al. Physical, mathematical, and numerical derivations of the Cahn–Hilliard equation. Computational Materials Science 81, 216–225 (2014). https://doi.org/10.1016/j.commatsci.2013.08.027

35 Sneider, A. et al. Deep learning identification of stiffness markers in breast cancer. Biomaterials 285, 121540 (2022). https://doi.org/10.1016/j.biomaterials.2022.121540

36 Shemesh, T., Geiger, B., Bershadsky, A. D. & Kozlov, M. M. Focal adhesions as mechanosensors: A physical mechanism. Proceedings of the National Academy of Sciences 102, 12383–12388 (2005). https://doi.org:10.1073/pnas.0500254102

37 Kim, D.-H. et al. Actin cap associated focal adhesions and their distinct role in cellular mechanosensing. Scientific Reports 2, 555 (2012). https://doi.org:10.1038/srep00555

38 Wu, Z., Plotnikov, S. V., Moalim, A. Y., Waterman, C. M. & Liu, J. Two Distinct Actin Networks Mediate Traction Oscillations to Confer Focal Adhesion Mechanosensing. Biophysical Journal 112, 780–794 (2017). https://doi.org/10.1016/j.bpj.2016.12.035

39 Gonnella, G., Marenduzzo, D., Suma, A. & Tiribocchi, A. Motility-induced phase separation and coarsening in active matter. Comptes Rendus Physique 16, 316–331 (2015). https://doi.org/10.1016/j.crhy.2015.05.001

40 Cates, M. E. & Tailleur, J. Motility-Induced Phase Separation. Annual Review of Condensed Matter Physics 6, 219–244 (2015). https://doi.org:10.1146/annurev-conmatphys-031214-014710

41 Maini, P. K., Woolley, T. E., Baker, R. E., Gaffney, E. A. & Lee, S. S. Turing’s model for biological pattern formation and the robustness problem. Interface Focus 2, 487–496 (2012). https://doi.org:10.1098/rsfs.2011.0113

42 Du, W., Nair, P., Johnston, A., Wu, P.-H. & Wirtz, D. Cell Trafficking at the Intersection of the Tumor–Immune Compartments. Annual Review of Biomedical Engineering 24, 275–305 (2022). https://doi.org:10.1146/annurev-bioeng-110320-110749

43 Jayatilaka, H. et al. Synergistic IL-6 and IL-8 paracrine signalling pathway infers a strategy to inhibit tumour cell migration. Nature Communications 8, 15584 (2017). https://doi.org:10.1038/ncomms15584

44 Stroka, Kimberly M. et al. Water Permeation Drives Tumor Cell Migration in Confined Microenvironments. Cell 157, 611–623 (2014). https://doi.org/10.1016/j.cell.2014.02.052

45 Bera, K. et al. Extracellular fluid viscosity enhances cell migration and cancer dissemination. Nature 611, 365–373 (2022). https://doi.org:10.1038/s41586-022-05394-6

46 Kasinrerk, W., Tokrasinwit, N. & Phunpae, P. CD147 monoclonal antibodies induce homotypic cell aggregation of monocytic cell line U937 via LFA-1/ICAM-1 pathway. Immunology 96, 184–192 (1999). https://doi.org:10.1046/j.1365-2567.1999.00653.x

47 Campanero, M. R. et al. An alternative leukocyte homotypic adhesion mechanism, LFA-1/ICAM-1-independent, triggered through the human VLA-4 integrin. J Cell Biol 110, 2157–2165 (1990). https://doi.org:10.1083/jcb.110.6.2157

48 Gilkes, D. M., Semenza, G. L. & Wirtz, D. Hypoxia and the extracellular matrix: drivers of tumour metastasis. Nature Reviews Cancer 14, 430–439 (2014). https://doi.org:10.1038/nrc3726

