## Supplementary Fig for "Mechano-induced homotypic patterned domain formation by monocytes"

### Supplementary Information

#### ***Detailed derivation of the Cahn-Hilliard equation for cell density***

The free energy density is composed of a bulk energy part ($\mathrm{X}$) and interfacial energy part ($I$):

$$f = \mathrm{X}(\rho) + I(\rho) = k_4(c_1\rho^4 + c_2\rho^3 + c_3\rho^2) + \frac{1}{2}k_3 e^{-\alpha c}(\nabla\rho)^2$$

The chemical potential $\mu$ is defined as the variational derivative of free energy with respect to the density $\rho$. To obtain the variational derivative, we firstly define the variation of density as: $\rho_\epsilon = \rho + \epsilon w$ where $\epsilon$ is a small perturbation. The variational derivative of the total free energy $G$ can then be written as:

$$\frac{d}{d\epsilon}\int_\Omega [\mathrm{X}(\rho+\epsilon w) + I(\rho+\epsilon w)]\, dV|_{\epsilon=0}$$

$\Omega$ is the whole region where cells live. This derivative can be further rewritten as:

$$\int_\Omega \left[\frac{d\mathrm{X}}{d\rho}\bigg|_{\rho_\epsilon} \frac{d}{d\epsilon}(\rho+\epsilon w) + (k_3 e^{-\alpha c}\nabla\rho)\cdot\nabla w\,\right] dV$$

$$= \int_\Omega [\mathrm{X}' w + (k_3 e^{-\alpha c}\nabla\rho)\cdot\nabla w] dV$$

Integrating by parts, we can get:

$$\frac{d}{d\epsilon}G[\rho_\epsilon]|_{\epsilon=0} = \int_\Omega \{\mathrm{X}' w - [\nabla\cdot(k_3 e^{-\alpha c}\nabla\rho)]w\} dV + \int_{\partial\Omega} w(k_3 e^{-\alpha c}\nabla\rho)\cdot\boldsymbol{n}\, dl$$

Where $\partial\Omega$ is the boundary of the region. The first term gives the chemical potential $\mu$:

$$\mu = \mathrm{X}' - [\nabla\cdot(k_3 e^{-\alpha c}\nabla\rho)]$$

At equilibrium state, $\mu = 0$ and $(k_3 e^{-\alpha c}\nabla\rho)\cdot\boldsymbol{n} = 0$ on boundary $\partial\Omega$. When the system is far from equilibrium, there will be flow: $\boldsymbol{j} = -D_1\nabla\mu$ and the governing equation for cell density can be written as: $\frac{\partial\rho}{\partial t} = -\nabla\cdot j = D_1\nabla^2\mu$. After substituting chemical potential into the diffusion equation, we can get:

$$\frac{\partial\rho}{\partial t} = D_1\nabla^2[k_4(4c_1\rho^3 + 3c_2\rho^2 + 2c_3\rho) - k_3 e^{-\alpha c}(-\alpha\nabla c\cdot\nabla\rho + \nabla^2\rho)]$$

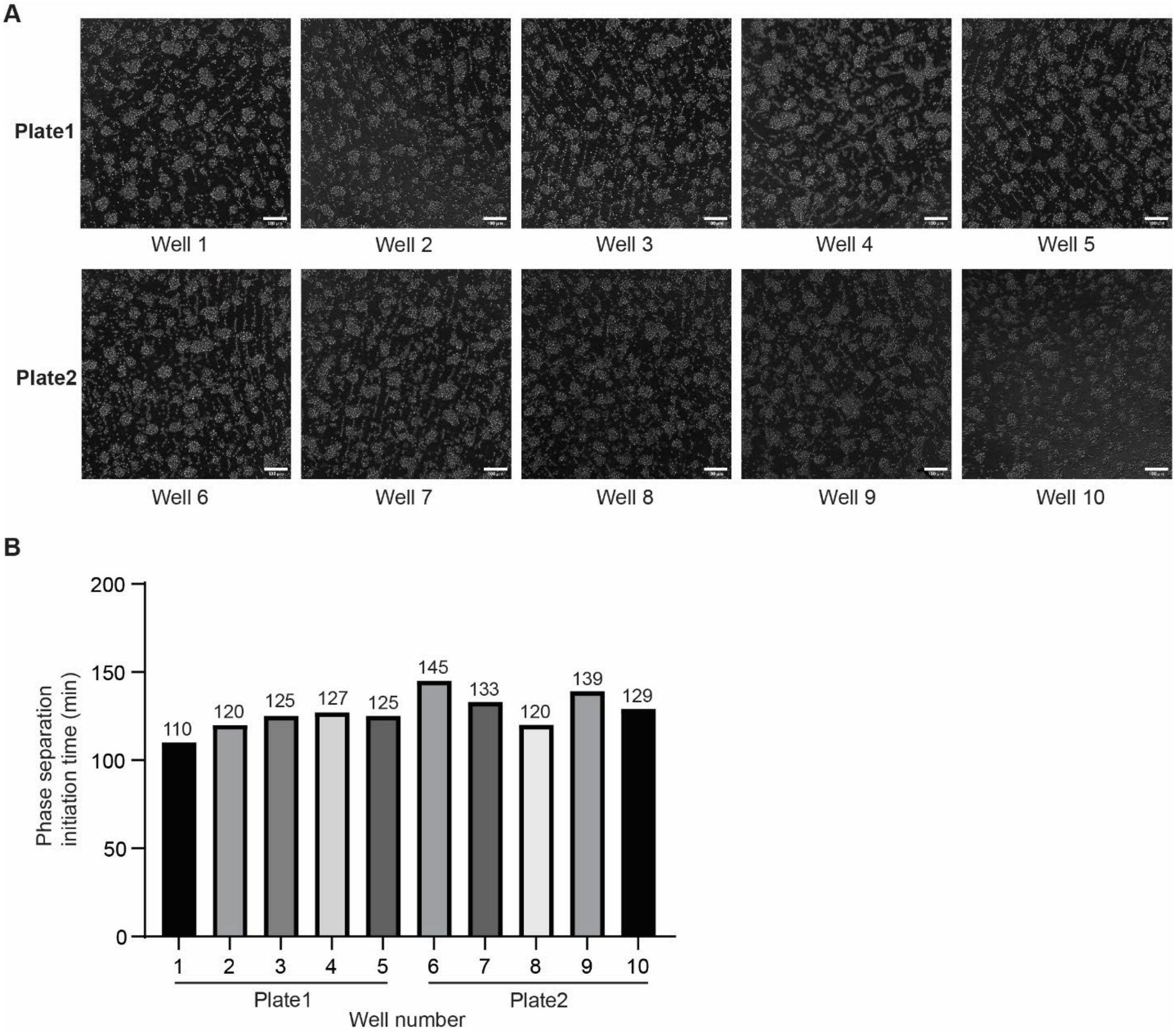


**Supplementary Fig. 1 Monocyte homotypic domain formation as a bio-physical phenomenon with great consistency. (A)** Representative 20x images of monocyte domain formations at 12h with primary monocytes isolated from same donor's PBMC seeded on different batches of commercially available 0.5 kPa collagen-coated polyacrylamide gels. Similar patterns were formed across all repeats. **(B)** great consistency existed in the phase separation initiation times for monocytes in repeats, which is around 120 minutes.

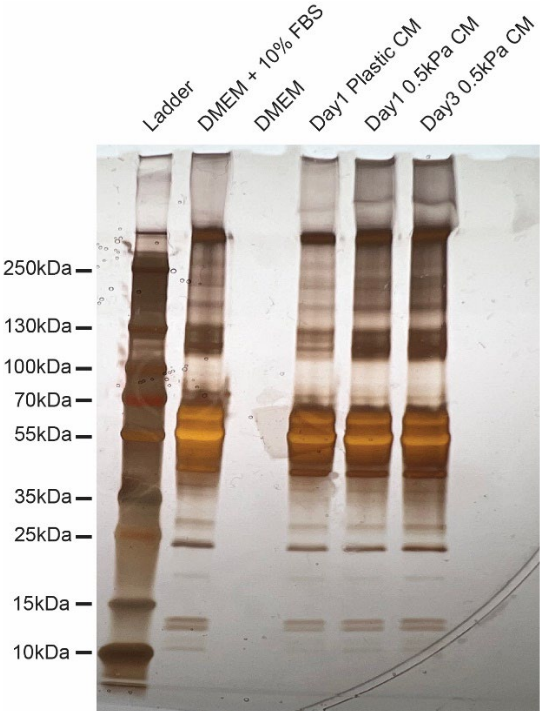


**Supplementary Fig. 2 Silver staining of concentrated conditioned media from "domained" monocytes on plastic and 0.5 kPa substrates.** Normalized to same number of cells, soluble factors around 110-130kDa were found to be differentially secreted by "domained" monocytes on 0.5 kPa physiological substrate compared to plastic substrate, which supports the assumption we made on the existence of inhibitory soluble factors in the model.

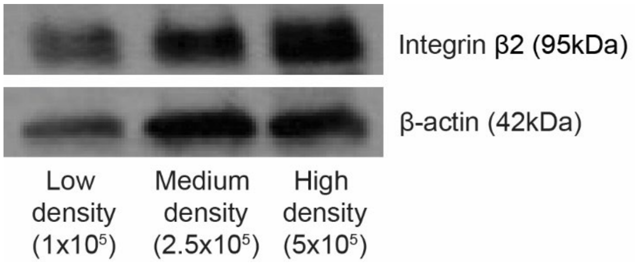


**Supplementary Fig. 3 Western Blot on integrin β2 expressions of monocytes seeded at low/medium/high density.** Elevated expression of integrin β2 was observed with increased seeding density.
